# Nasal inhalations and exhalations evoke distinct prefrontal-cortex responses with constant latency

**DOI:** 10.64898/2026.07.21.739551

**Authors:** Diego A. Laplagne, Joseph A. Belo, Ana L. A. Dias, Elis H. Duarte, Alan M. B. Furtunato, Adriano B. L. Tort

**Author notes:** Equal contribution (Diego A. Laplagne, Joseph A. Belo).

## Abstract

Neuronal activity synchronous to breathing is salient across the mammalian brain. These modulations are typically characterized as oscillating signals phase-locked to the ongoing breathing rhythm and treated as evidence for respiratory ‘entrainment’ of cortical and subcortical oscillations. Since activation of the olfactory circuitry has been identified as their preponderant source, we wondered whether respiratory brain signals could be better understood as sequences of evoked sensory potentials. To help distinguish between these scenarios, we set out to quantify the detailed alignment of prefrontal cortex local field potentials (LFPs) to the nasal airflow cycle across the natural repertoire of rat breathing. We found that, while LFP and breathing show salient synchrony for all breathing modes, the phase of their locking is a strict linear function of respiratory rate. We explain this by showing that LFPs align to nasal airflow with a constant time lag of 100 ms, irrespective of breathing rate. Segmenting by breathing cycles revealed that LFP peaks and fast-gamma bursts are time-locked to inhalations, while LFP troughs and slow-gamma activity time-lock to exhalations. Our results support the view that respiratory signals across the frontal brain of rodents are sensory potentials evoked by cyclic ortho- and retronasal airflow.

## Introduction

Animals continuously collect information about the external and internal world. There is renewed interest in understanding how interoceptive signals reach the central nervous system and interact with exteroceptive sensory inputs to shape behavior, emotion and cognition [1,2]. Dissecting the circuits and mechanisms through which bodily signals impact neuronal activity across brain regions is key to unveiling their functions [3]. Mammalian breathing is a versatile motor pattern that flexibly balances the need for respiration with behavioral demands like vocalizing, sniffing and swallowing [4]. Remarkably, recent work has shown that respiratory signals are globally broadcast, as breathing strongly modulates neuronal activity across the brain of mammals — humans included [5]. At the prefrontal cortex (PFC), neurons, assemblies, field potentials and gamma oscillations show robust respiratory synchrony [6–19]. The functional roles played by brain-breathing signals are a matter of active research, with possible contributions highlighted for sensory coding [20–23], perception and action [24–27], emotional coordination [6,9,15,28,29], and memory [30–33].

Multiple lines of evidence have established sensory input from the nose as a preponderant source for respiratory neural modulations across the brain [33–36], specifically via the activation of the olfactory circuit [6,7,16,37–41]. During inhalations, low pressure in the airways creates orthonasal (inwards) airflow carrying odorants from the exterior while, during exhalations, high pressure produces retronasal (outwards) flow of air enriched in volatile odorants from the mouth and digestive system [42]. This results in the cyclic activation of olfactory sensory neurons (OSNs) in the olfactory epithelium (OE) [43,44].

Notably, OSNs are themselves mechanosensitive, such that they are directly activated by both odorants and air pressure in the nasal cavity [45–47]. OE activity propagates via the olfactory bulb (OB) directly to olfactory and entorhinal cortices and cortical amygdala [48–51], and indirectly to structures across the brain, including the hippocampus, thalamus and PFC [52–55], all of which were shown to be modulated by the breathing rhythm [7,11,13,16,17,33–37,56,57].

The alignment between oscillatory signals can be measured with either frequency- or time-domain analysis tools, respectively quantifying the phase difference or latency between them [58]. For signals with stable frequency, constant phase implies constant latency and vice versa, but this does not hold when the oscillation frequency varies (e.g. a latency of 0.25 s corresponds to a phase delay of a quarter-cycle for 1-Hz signals but becomes a half-cycle for 2 Hz). The respiratory modulation of sensory coding in the OB has been successfully studied by aligning neuronal activity to the ongoing phase of the nasal breathing cycle in controlled settings with stable breathing rates [21,59]. Interestingly, recent analyses have unveiled OB neural signals with constant latency to the breathing pattern across a wider range of rates [12,60–62], in line with the fixed stimulus-response delays characteristic of sensory systems [63–65].

Neuronal activity in higher brain circuits is paced by salient endogenous oscillations [66–68], and respiratory modulation of cortical and subcortical activity is commonly studied as oscillatory ‘entrainment’ of these oscillations by the breathing rhythm. Entrainment occurs when two oscillators, which could follow different frequencies when isolated (their ‘natural’ frequencies), become phase-coupled through unidirectional or bidirectional coupling [69,70]. Determining whether a periodic sensory input entrains or evokes brain activity can be challenging, as both mechanisms produce activity that is frequency-locked to the rhythm of the inputs [71]. Quantifying the alignment between sensory input and brain signals across a wide range of rates can help disambiguate these two mechanisms [72]: while fixed latencies are expected for feedforward synaptic broadcasting of sensory activity, stable phase differences across rates would support oscillatory entrainment [72–74]. Using frequency-analysis tools, global respiratory brain rhythms are characterized under the entrainment paradigm as neural signals oscillating at the ongoing breathing rate with significant phase-locking to it [11,16,36,56,75]. Alternatively, some studies have shown brain signals time-locked to breathing [33,38,76]. Crucially, if and how the phase difference or latency between brain activity and breathing depend on respiratory rate has not been studied for high-order areas.

In this work, we recorded nasal breathing and distributed brain potentials in the OB and PFC of rats as they freely transitioned between cognitive and behavioral states. We then combined novel and established analysis techniques to quantify the phase and time delay between neural and respiratory signals across the wide range of rat breathing rates. We show that frontal brain activity harbors specific low-and high-frequency components aligned with fixed latency to nasal inhalations and exhalations, supporting the view that respiratory signals across the frontal brain of rodents are sensory potentials evoked by cyclic ortho-and retronasal airflow.

## Methods

### Animal subjects and recording sessions

All experimental procedures were approved by the Ethics Committee for Animal Experimentation of the Federal University of Rio Grande do Norte (CEUA, protocol no. 062.070/2017), in accordance with Brazilian Federal Law No. 11.794/2008 (Lei Arouca).

Animals used in this study were the same 7 male Wistar rats (3 months old) as in [28]. Rats were stereotaxically implanted with a custom-made 14-electrode array (2 parallel rows of 7 tungsten wires, 75 μm, 500-μm spacing; see Figure S3a) spanning the medial aspect of the left prefrontal cortex (PFC), targeting the medial-orbital (MO), prelimbic (PRL) and anterior cingulate (ACG) areas (AP, 1.2–4.2 mm; ML, 0.4–0.9 mm; DL, −3.5 mm for MO). The olfactory bulb (OB) was recorded from a stainless steel screw over its right dorsal surface (AP, 7.5 mm; ML, −1 mm) and a second screw over the left parietal cortex (AP, −4 mm; ML, +3 mm) served as ground and reference. Breathing was monitored through a stainless steel cannula implanted through the nasal bone into the right nasal cavity, reversibly connected to a head-mounted pressure sensor (piezoresistive bridge; < 1 ms response time) during recordings. All signals were acquired at 12,207 Hz with a wired headstage. Experiments were conducted at the beginning of the dark phase of a 12 h light/dark cycle. See [28] for additional detail and histology.

Rats were recorded in the dark as they freely explored a square arena of 0.7 m side length and 0.5 m height. The arena floor was covered in wood shavings with two plastic objects placed as enrichment, one of them containing soiled bedding from a cage housing a female rat. Before recording, the rats were habituated to the human experimenter, but not to the arena. We monitored the recordings through infrared-based depth video (Kinect2, Microsoft). We recorded 2 to 5 sessions per rat, amounting to a mean valid recording time of 5231 s per rat (range 2691 to 7159 s; 10.2 hours of total recording time). These open-field sessions were interleaved with the elevated-plus maze sessions in [28] and sessions in a social arena, leaving 2-day rest periods between recordings at each arena.

### Data analysis

All analyses were conducted with custom-made routines in MATLAB R2018a (The Mathworks). LFP and nasal-pressure signals were anti-aliased at 500 Hz (zero-phase Butterworth, order 3) before downsampling to 1220.7 Hz. Signals were concatenated over all sessions for each rat. LFP electrodes with visible signal artifacts, and time periods with poor pressure or LFP signal quality were discarded. We inverted the nasal pressure signal (inhalations up) throughout and the OB signal for the analyses in Figures 1 and 2. The OB signal was lost for one rat, leaving an *n* of 6 for all OB analyses.

**Figure 1.**
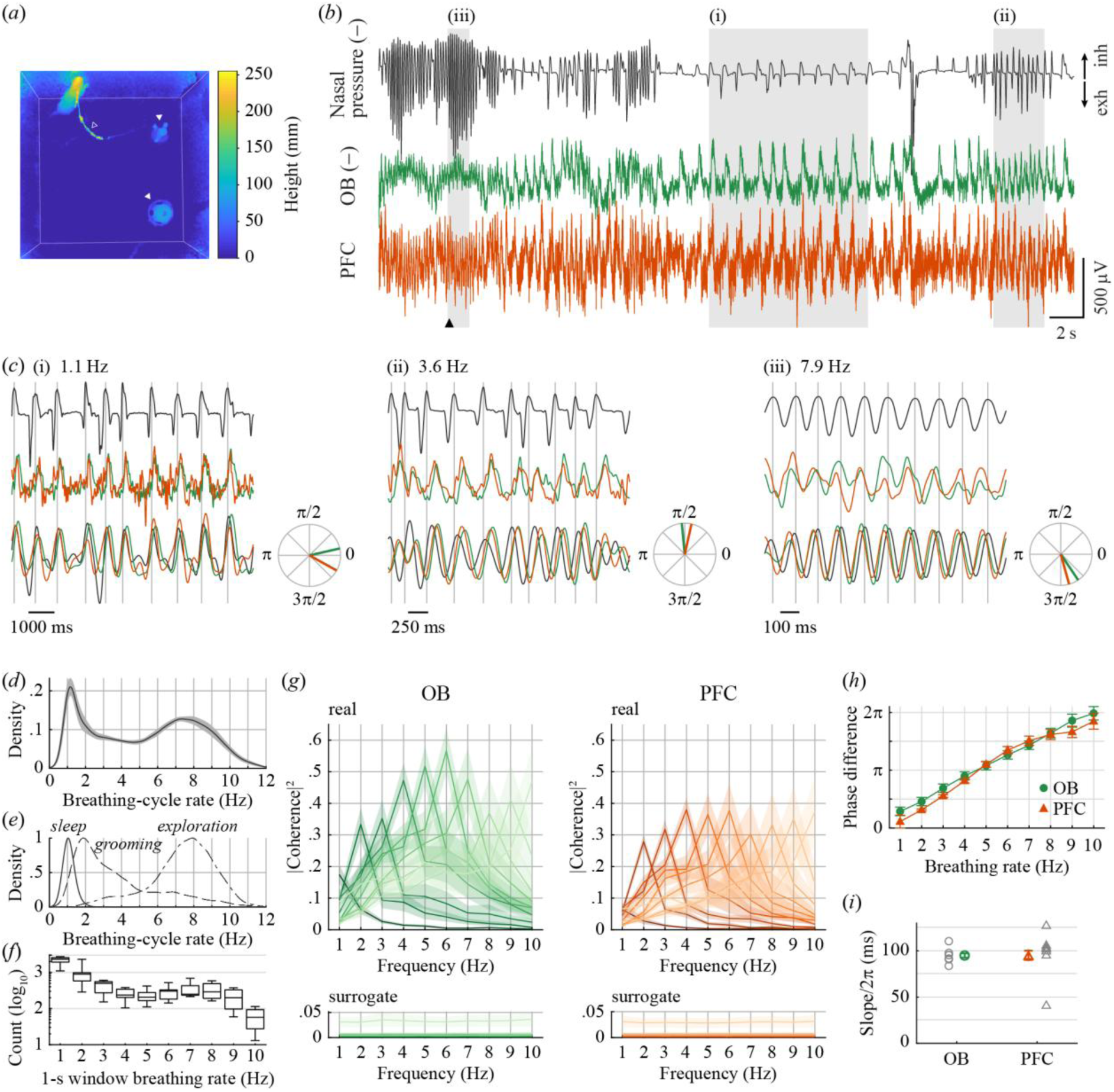
Phase difference between frontal brain potentials and nasal breathing across respiratory rates. **(*a*)** Top view of the open-field arena (70 × 70 cm), as recorded with a depth camera (color is height from the arena floor). A rearing rat can be seen, with its headstage cable (open triangle), as well as two objects on the floor (closed triangles). White lines mark the edges of the arena. **(*b*)** Example traces of intranasal pressure (top) and OB (middle, green) and PFC (MO electrode; bottom, orange) LFPs, with pressure and OB traces inverted (here and throughout all figures, see text). Pressure and LFPs are filtered at .1–20 Hz and .1–100 Hz respectively, and pressure is scaled in arbitrary units. Grayed rectangles mark the time periods expanded in (*c*) and arrowhead marks the time of the video frame in (*a*). **(*c*)** Detailed views of the time periods highlighted in (*b*), each comprising 10 respiratory cycles, with mean breathing-cycle rates of 1.1 Hz (i), 3.6 Hz (ii) and 7.9 Hz (iii). Top: nasal pressure; Middle: OB (green) and PFC (orange) LFPs, filtered at .1–20 Hz; Bottom: nasal pressure and LFPs, narrow-band filtered at breathing rate ± 1 Hz. Polar plots: mean phase difference from OB and PFC LFPs to nasal pressure for the narrow-band-filtered signals. **(*d*)** Distribution of breathing-cycle rates (kernel-density estimation; bandwidth 0.25 Hz). **(*e*)**Distribution of breathing-cycle rates for periods of NREM sleep (solid), grooming (dashed) and exploratory behaviors (dot-dashed). **(*f*)** Number of 1-second windows at each 1-Hz rate bin (median, [.25, .75] quantiles and range over rats). **(*g*)** Mean magnitude-squared coherence spectra between nasal pressure and LFPs from the OB and PFC for 1-second data windows sorted in bins by their peak breathing rate from 1 Hz (darkest) to 10 Hz (lightest). Note that each spectrum peaks at the corresponding breathing rate. Spectra from window-shuffling surrogate controls are depicted below (note the difference in vertical scales). **(*h*)** Phase of breathing-brain coherence for OB and PFC as a function of breathing rate. **(*i*)** Predicted time-lags from nasal pressure to OB and PFC LFPs, as estimated from dividing phase-rate slopes by 2· π (positive values correspond to LFP trailing nasal pressure). Values in (*d*, *g*–*i*) are mean ± sem over rats; gray markers in (*i*) are for individual rats; (*e*) is mean over rats, normalized to maximum density for clarity.

**Figure 2.**
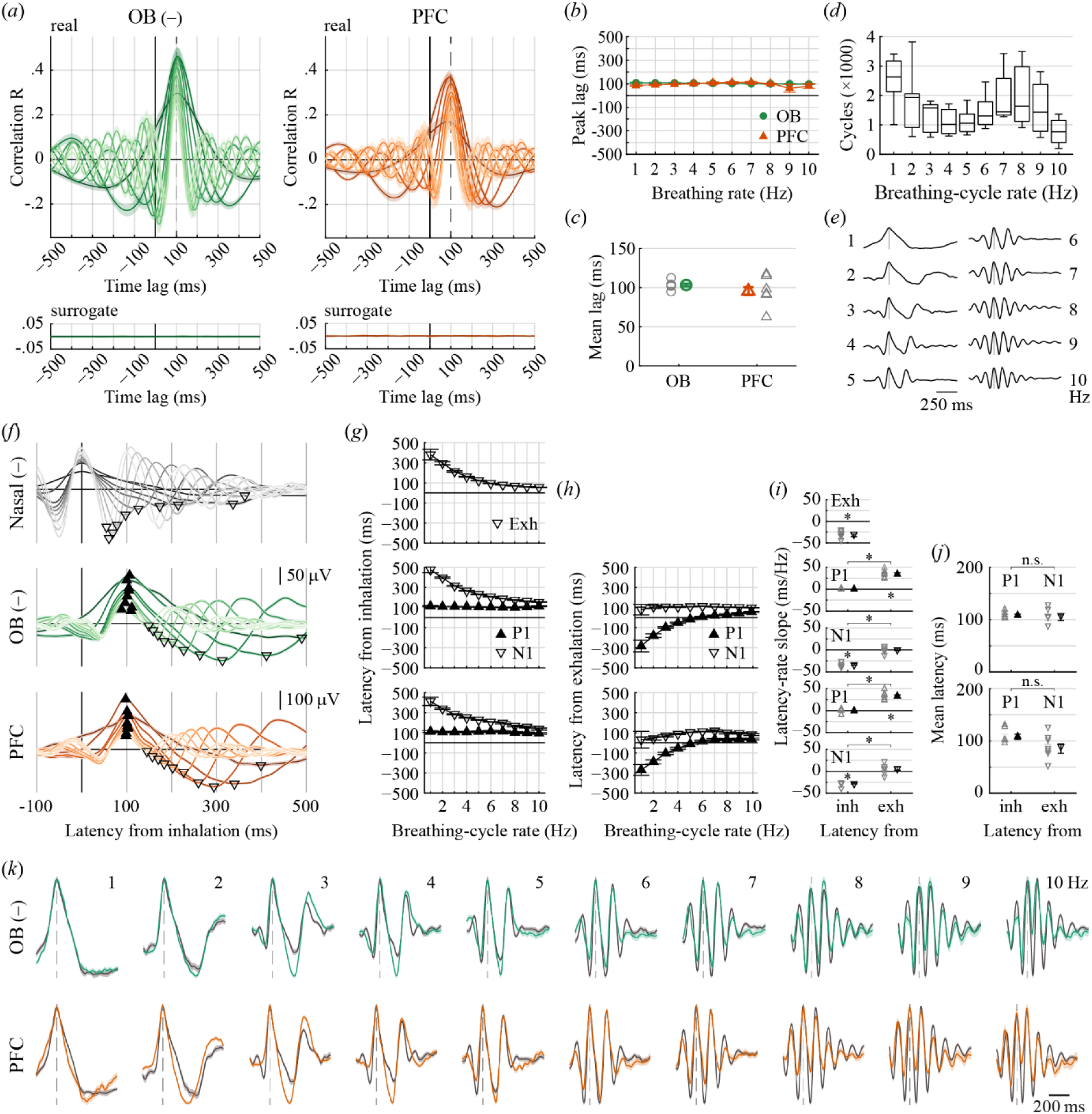
Latency between frontal brain potentials and nasal breathing across respiratory rates. **(*a*)** Mean cross-correlograms between nasal pressure and LFPs from the OB and PFC for 1-second data windows sorted in bins by their peak breathing rate from 1 Hz (darkest) to 10 Hz (lightest); dashed line marks +100 ms lag (positive time-lag values correspond to LFP trailing nasal pressure). Cross-correlograms from window-shuffling surrogate controls are depicted below (note the difference in vertical scales). **(*b*)** Lag at LFP-breathing cross-correlation peaks for OB and PFC as a function of breathing rate. **(*c*)** Mean over breathing rates of cross-correlation-peak lags. **(*d*)** Number of breathing cycles at each 1-Hz rate bin (median, [.25, .75] quantiles and range over rats). **(*e*)** Inhalation-triggered averages (ITAs) of nasal pressure for breathing cycles in each 1-Hz rate bin for an example rat, normalized by inhalation peak amplitude (bootstrap mean ± sem; gray line marks the time of peak inhalation). **(*f*)** ITAs of nasal pressure (top) and OB and PFC LFPs (middle and bottom) for breathing cycles in each rate bin, superimposed and colored by rate from darkest (1 Hz) to lightest (10 Hz; same example rat as in (*e*), bootstrap mean ± sem). For nasal pressure, open triangles mark peak exhalations for each rate. For LFPs, closed and open triangles mark the first post-inhalation positive (“P1”) and post-exhalation negative (“N1”) voltage peaks for each rate. **(*g*)** Latency of peak exhalation (top) and P1 and N1 (middle and bottom), measured from times of peak inhalation, as a function of breathing-cycle rate. **(*h*)** Latency of P1 and N1, measured from times of peak exhalation, as a function of breathing-cycle rate. **(*i*)** Linear slopes of latency vs. breathing-cycle rate. Top: slopes for latencies of exhalation peaks from inhalations; Middle and bottom: slopes for latencies of P1 and N1 measured from inhalations (“inh”) and exhalations (“exh”). **(*j*)** Mean over breathing-cycle rates of latencies of P1 from inhalations and N1 from exhalations. **(*k*)** Superimposed ITAs of nasal pressure (gray, delayed by 100 ms) and OB (green, top) or PFC (orange, bottom) LFPs, for breathing-cycle rate bins of 1 to 10 Hz (left to right). Same traces as in (*f*), normalized by inhalation or P1 amplitude; dashed lines mark inhalation peaks. * significant (all p < 0.0005); n.s.: not significant. Values in (*a*– *c*, *g*–*j*) are mean ± sem over rats; gray markers in (*c*, *i*, *j*) are for individual rats.

For each rat, we selected the cortical electrode with the largest breathing modulation for analysis (‘PFC’). This allowed us to maximize signal-to-noise when quantifying the alignment of LFP signals to breathing. Specifically, we selected the channel with the largest inhalation-triggered average RMS at 0–500 ms latencies for breathing cycles in the 1.5–2.5 Hz range. This electrode was located in the MO for 5 rats and in the PRL for the other 2. We reproduced selected analyses across cortical areas by first obtaining one mean LFP signal per area for each rat (Figure S1).

For the analysis of current flow density (Figures S3 and S6), we obtained differential LFPs (∇LFP) between pairs of cortical electrodes (0.5 mm distance; 2 pairs in the MO, 3 in the PRL and 2 in the ACG) by subtracting the signals in the lateral electrode from that of the medial one and dividing by the electrode distance; positive ∇LFP reflects positive current flow from the medial to the lateral electrode. Only pairs where both electrodes had valid signals were included from each rat (see Supplementary Methods).

#### Breathing

Nasal pressure was demeaned and high-pass filtered at 0.1 Hz (zero-phase Butterworth, order 3) to remove any baseline fluctuations. We inverted the pressure signal, leaving inhalations up. For peak detection, the signal was then smoothed via convolution with a Gaussian kernel of full-width at half maximum of 25 ms and inhalation peaks were identified with the built-in *findpeaks* function with MinPeakDistance at 1/12 (maximum cycle rate of 12 Hz; we define each breathing cycle as the period between consecutive inhalation peaks, and its rate as the inverse of its duration). The accuracy of peak detection was visually confirmed for all recordings.

For Figure 1e, periods of grooming (269 to 1776 s and 682 to 4634 breathing cycles per rat) and exploration (walking, object exploration or rearing; 179 to 300 s; 882 to 2220 cycles) were manually staged from the depth videos. NREM (462 to 1089 s; 411 to 1246 cycles) was defined as immobility with high delta (0.75–4 Hz) power in a PRL electrode.

#### Breathing-brain coherence

We developed an adaptation of coherence analysis for quantifying its amplitude and phase at the time-varying instantaneous rate of one of the two signals (*rate-resolved coherence*; C_RR_). For the case of breathing-LFP coherence, the rate of interest is the fundamental frequency of the respiratory signal, that is, the ongoing breathing rate. As detailed below, we effectively segmented the data for each rat into 1-second windows, sorted them into 1–10 Hz breathing-rate bins and calculated the coherence magnitude and phase over each pool of windows at the frequency matching its respiratory rate.

Our method begins by segmenting the nasal-pressure (inverted) and LFP (OB inverted) signals in consecutive, non-overlapping windows of 1-second duration and removing the mean from each windowed signal. We then obtain the single-sided Fourier spectrum (built-in *fft* function) of each window from each signal and crop it to our frequency range of interest (1–10 Hz, 1-Hz resolution). For each window, we obtain the breathing and LFP power spectra by elementwise multiplying each Fourier spectrum by its complex conjugate, and their cross-spectrum by elementwise multiplying the breathing spectrum with the complex conjugate of the LFP spectrum. The critical step comes next, where we estimate the breathing rate of each time window as the frequency with maximum power in the breathing signal and then sort all the spectra above into 1-Hz breathing-rate bins, from 1 to 10 Hz.

For each breathing-rate bin, we then computed 1000 bootstrap estimates of the complex coherency spectrum, each from *n* windows sampled with replacement, where *n* is the total number of windows in the bin. Each estimate is the mean breathing-LFP cross-spectrum divided by the square root of the product of the mean breathing and LFP power spectra:

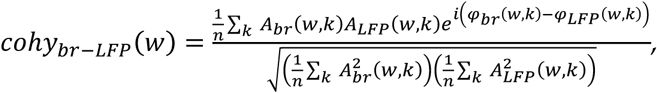

where *w* is frequency, *k* is window number, *A* is Fourier spectrum amplitude and φ is phase. From each coherency estimate, we computed the magnitude-squared coherence (MSC) spectrum as its squared absolute value and the phase spectrum as its angle. The MSC and phase spectra of each rate bin are, respectively, the mean and the circular mean of its 1000 bootstrap estimates (MSC spectra are shown in Figure 1g).

We constructed the C_RR_-magnitude and C_RR_-phase functions by taking, for each rate bin, the MSC and phase values at the frequency matching the center of the bin; we applied the built-in *unwrap* function to the rate-phase curve of each rat (C_RR_-phase functions are shown in Figure 1h). Phase difference is in reference to the breathing signal (i.e.: a value of *π*/2 represents nasal pressure leading the LFP by a quarter-cycle). We assessed the significance of the C_RR_ magnitude for each combination of rat and rate by contrasting its 1000 bootstrap estimates against 1000 surrogate estimates from within-rate window-shuffling.

#### Cross-correlation

We analyzed cross-correlation between the nasal pressure and LFP signals for each rat using the same 1-second data windows sorted in breathing-rate bins described above. We computed 1000 bootstrap estimates of the mean cross-correlogram for each rate bin, normalized to Pearson correlation R values, and obtained the lag and R for the maximum of each estimate. We then kept the mean curve (Figure 2a), mean peak lag (Figure 2b) and mean peak R over bootstrap estimates for each rate bin. We assessed the significance of peak R by contrasting the 1000 bootstrap estimate values against 1000 surrogate estimates from within-rate window-shuffling.

#### Inhalation-triggered average (ITA)

We implemented ITAs as event-triggered averages of the nasal-pressure or LFP signals, with times of inhalation peaks as event times (Figure 2f). The events were sorted into 1-Hz bins by their corresponding breathing-cycle rate (the inverse of cycle duration). For each group of events, we extracted the signal windows at −0.5–1.5 s lag and obtained 1000 estimates of their mean waveform from subsamples of 500 events, sampled with replacement (fixing the sample size in this way controls for positive finite-sample bias when obtaining RMS values from the average waveform). From these, we computed the mean waveform and its standard error (sem) as the standard deviation over estimates. We assessed the significance of each ITA RMS in the 0–0.5 s lag interval by contrasting the value from the 1000 estimates against 1000 surrogate estimates obtained from random circular-shifts of each window over its full 2-second duration. We subtracted the grand mean of these surrogate waveforms from the real ITA waveform.

We used *findpeaks* to detect peaks on the ITA waveforms (Figure 2f–h). For exhalation peak (nasal pressure) and P1 (LFP), peak detection started at 40 ms after inhalation-peak time to avoid possible residual fluctuations from signals linked to the previous breathing cycle. For each breathing-cycle rate bin, we further set the MinPeakDistance parameter to 80% of its upper cycle-duration limit and kept the first positive peak detected. The selected peak was thus the most prominent within a lag window of approximately one breathing cycle. N1 was detected as the first negative peak after P1, running *findpeaks* on the inverted LFP signal with MinPeakDistance set as before. Latencies from exhalation were calculated by subtracting the exhalation-peak latency from the N1 latency for each rate bin.

#### Gamma bursts

For analyses in the gamma band, we selected the PFC representative channel for each rat as before for LFP, but based on the 50–100 Hz envelope (for only one rat this channel differed from the one selected for LFP, from one MO electrode to another). LFP spectrograms in the gamma band (30–110 Hz; Figure 3a, b) were power spectral density computed with the built-in *spectrogram* function with a window of 75 ms, 15/16 overlap and frequency resolution of 0.25 Hz. For each breathing-cycle rate bin, we obtained ITAs of the gamma spectrograms as the mean of 50 subsampling estimates (each of 500 events selected with replacement, −0.2–0.6 s lag; Figure 3c, d). We subtracted the surrogate mean (random circular-shift over each 0.8-second window) from the spectrogram ITAs.

**Figure 3.**
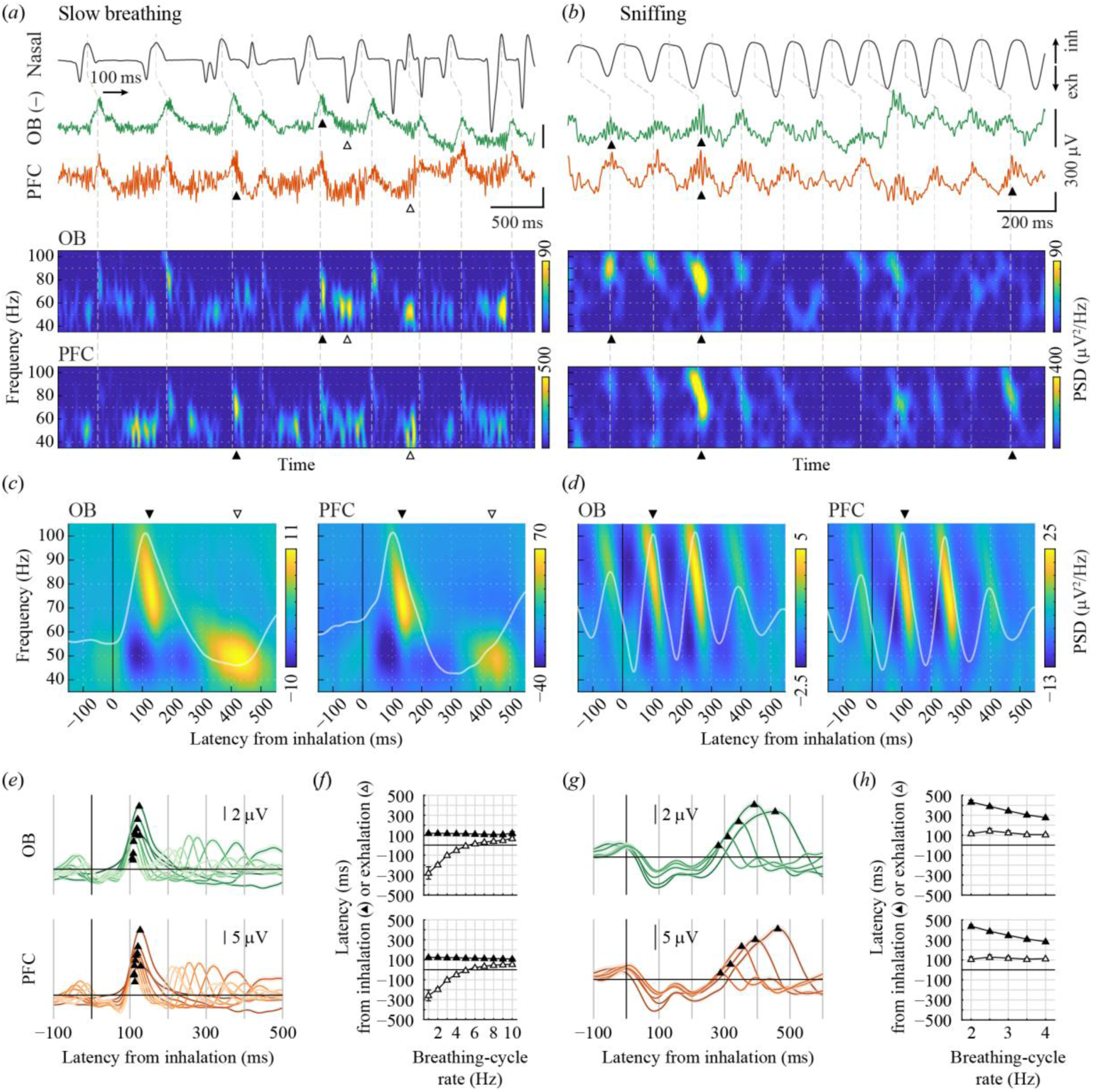
Latency between gamma oscillation bursts and nasal breathing across respiratory rates. **(*a*)** Top: Example traces of intranasal pressure (.1–20 Hz; top, black) and OB (middle, green) and PFC (MO electrode; bottom, orange) LFPs (.1–100 Hz) during a period of slow breathing (mean breathing-cycle rate 2.6 Hz). Dashed lines connect each inhalation peak to the point 100 ms later in the LFPs. Bottom: spectrograms of the LFP signals. Closed and open arrowheads highlight prominent fast- and slow-gamma bursts, respectively. **(*b*)** As in (*a*), for a period of sniffing (mean breathing-cycle rate 7.4 Hz). **(*c*)** ITA of the gamma spectrogram for all breathing cycles in the 1.5– 2.5 Hz rate range from the same rat in *a, b* (870 cycles). PSD values represent the difference to the average surrogate spectrogram for the same cycles (see Methods). The waveforms of the corresponding LFP ITAs are superimposed in white for reference. Arrowheads as in (*a*) and (*b*). **(*d*)** As in (*c*), for breathing cycles in the 6.5–7.5 Hz rate range (1441 cycles). **(*e*)** ITAs of the OB and PFC fast-gamma envelope (70–90 Hz) for breathing cycles in 1-Hz rate bins, superimposed and colored by rate from darkest (1 Hz) to lightest (10 Hz; same example rat as in (*a*–*d*), bootstrap mean ± sem). Triangles mark the detected gamma envelope peak for each ITA. **(*f*)** Latency of fast-gamma envelope peaks, measured from times of inhalation (closed) or exhalation (open) peaks, as a function of breathing-cycle rate. **(*g*, *h*)** As in (*e, f*), for slow-gamma envelopes (50–55 Hz; 0.5-Hz bins). Values in (*f*, *h*) are mean ± sem over rats.

To quantify the latencies of the fast- and slow-gamma bursts to breathing-cycle landmarks (Figure 3e–h), we first band-pass filtered the LFP signals with *eegfilt* from the EEGLAB toolbox [77], within 70–90 Hz and 50–55 Hz for fast and slow gamma, respectively. From these, we computed the gamma envelopes by rectifying and low-pass filtering them to 25 Hz. We obtained the ITAs of the gamma envelopes, detected their first positive peaks and computed their latencies from inhalation and exhalation peaks as described above for the P1 peaks.

For the cycle-by-cycle correlations of gamma amplitude (Figure 4), we obtained the gamma envelopes as before and independently z-scored them. For each breathing cycle and electrode, we then obtained the peak envelope amplitudes within fixed latency windows (100–150 ms from inhalation for γ_fast_ and 50–150 ms from exhalation for γ_slow_; the latter was wider to account for larger inter-cycle variability). We conditioned on the predictor region (OB for OB-PFC correlations and PFC for PFC-PFC’ correlations), keeping only data from cycles with gamma amplitude > 2 z in it. For each pool of cycles reported (median 608 cycles, range 228–1876 per pool) we computed R^2^ and p-values from Pearson correlations. For gamma coherence we extracted the raw LFP signals from the same latency windows, detrended and multiplied each by a Hann taper and obtained MSC spectra with the formulations above over the cycles in each pool. From these we extracted MSC for γ_fast_ and γ_slow_ as the value at the frequency closest to the center of each gamma band (79 and 50 Hz).

**Figure 4.**
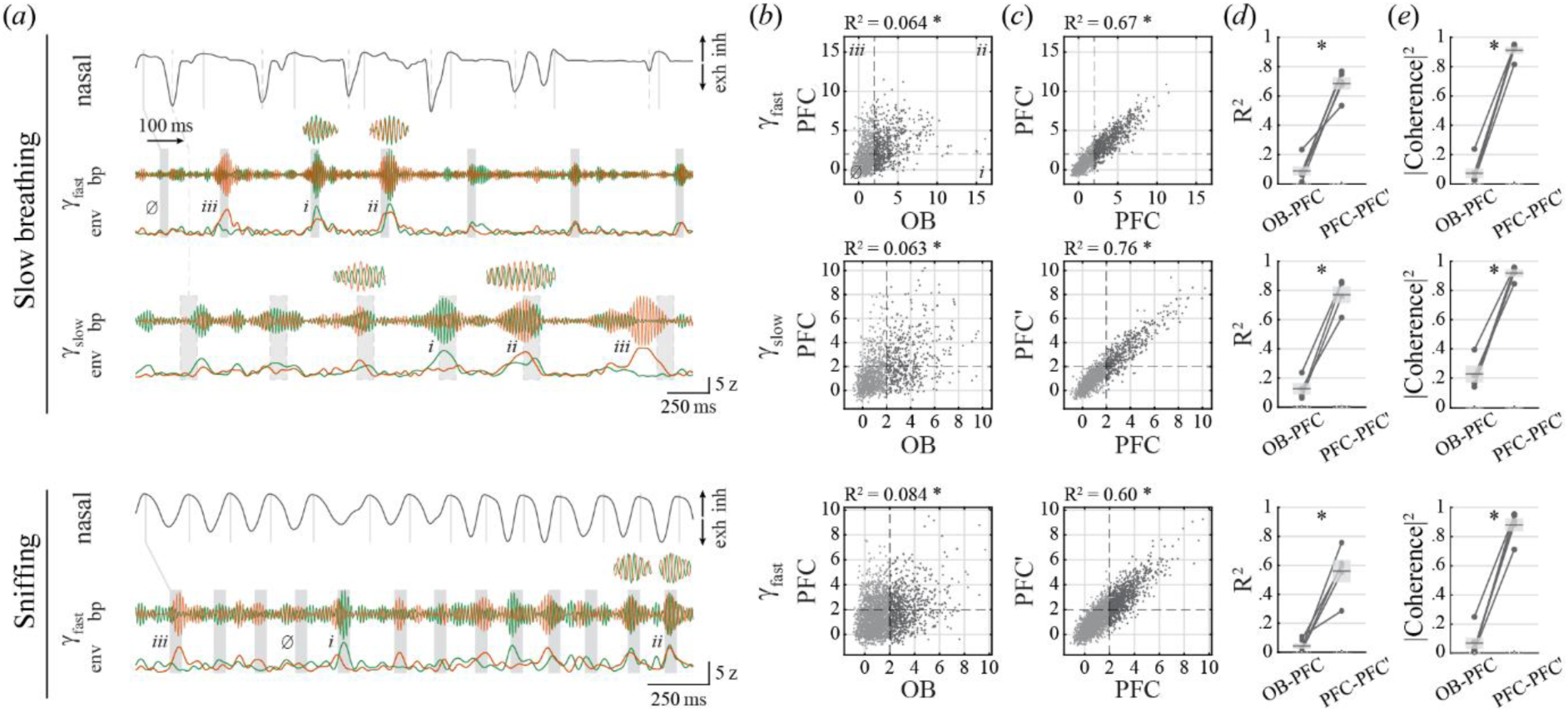
Cycle-by-cycle modulation of respiratory gamma bursts across the OB and PFC. **(*a*)** Top: Example traces of intranasal pressure (.1–20 Hz, black) and gamma-band LFPs at the OB (green) and PFC (MO electrode; orange) during a period of slow, irregular breathing (mean breathing-cycle rate 3.5 Hz). For each of fast and slow gamma, ‘bp’ is the band-passed signal (γ_fast_: 70–90 Hz; γ_slow_: 50–55 Hz; z-scored) and ‘env’ its envelope (z-scored). Solid and dashed lines indicate times of inhalation and exhalation peaks, respectively (two fast cycles not highlighted for clarity). Shaded rectangles mark the time windows for detecting gamma bursts in each cycle (γ_fast_: 100–150 ms from inhalation; γ_slow_: 50–150 ms from exhalation). Note example cycles for which gamma bursts were largest for OB (*i*), similar for OB and PFC (*ii*), largest for PFC (*iii*) or absent in both (∅). Insets: selected gamma bursts stretched 2× in time to expose their OB-PFC phase relationship. Bottom: As in top, for a period of sniffing (mean breathing-cycle rate 6.0 Hz). Respiratory γ_slow_ is not evident for fast breathing and was not analyzed. **(*b*)** Paired distributions of gamma burst amplitude (z-score units) in OB and PFC from one example rat, with each dot representing values from one breathing cycle; dashed lines mark 2-z boundaries and quadrant labels match the descriptions in (*a*). Throughout (*b–e*), slow breathing (top and middle rows) is cycles in the 2–4 Hz range and sniffing (bottom row) is cycles in the 6–8 Hz range. Analyses in (*d*, *e*) include only cycles with OB gamma > 2 z for OB-PFC and PFC gamma > 2 z for PFC-PFC’. **(*c*)** As in (*b*), for amplitudes in the PFC channel vs. a second channel in the PFC (PFC’). **(*d*)** R^2^ coefficients for OB-PFC and PFC-PFC’ linear correlations, rat by rat. **(*e*)** Magnitude-squared coherence values for OB-PFC and PFC-PFC’ pairs, rat by rat. * are p < 1e^−6^ for (*b*, *c*) and p ≤ 0.005 for (*d*, *e*). In (*d*, *e*), horizontal lines and shaded boxes are mean ± sem over rats (solid lines for real values, dashed lines for surrogates; note surrogates are at near-zero values).

For contrasts between pairs of PFC signals, we selected a second cortical channel (PFC’) as that with the second largest significant breathing modulation of gamma for each rat (one rat had no second significant channel and was not included in PFC-PFC’). For both correlation and coherence, surrogate values were obtained as the means of 1000 cycle-shuffles for each pool.

#### Statistics

Linear mixed models (LMM) were fitted with the built-in *fitlme* function using restricted maximum likelihood (REML). For models spanning rats, breathing rates and brain regions (as in Figure 1h), we included fixed effects for brain region, breathing rate (centered by subtracting the range midpoint, 5.5 Hz) and their interaction, plus a random intercept per rat. Slopes and midpoint-rate values reported for these models are grand marginal estimates. R^2^ values are conditional (fixed + random effects) and adjusted for the number of fixed-effects coefficients. *F* and p-values for the fixed effects were computed with the built-in *anova* function; *F* values are for the reference region (OB or MO). For models with 3 brain regions (as in Figure S1), p-values for post-hoc contrasts between regions were corrected with the Holm-Bonferroni method. Breathing-brain coherence phase and cross-correlation peak lag values from combinations of rat, brain region and breathing rate not significant at α = 0.01 were excluded from their respective LMMs. Linear models independently constructed for each rat and brain region combination (as for the slope values in Figure 1i) were fitted with the built-in *fitlm* function. For contrasts against surrogate distributions, we obtained one-sided p-values with finite-bias correction [78]. All p-values smaller than 1e^−6^ are reported as p < 1e^−6^.

## Results

We recorded breathing and frontal brain activity from rats freely behaving in an enriched open field in the dark (Figure 1a). We monitored breathing via intranasal pressure, whose high signal-to-noise ratio and frequency response allow for the reliable detection of individual breathing cycles with close-to-zero phase lag across the physiological range of rat breathing rates [79,80]. Nasal pressure is directly proportional to airflow [81], with peak positive and negative pressure coinciding with peak outflow and inflow during exhalation and inhalation, respectively. We simultaneously recorded local field potentials (LFP) from electrodes in the dorsal surface of the OB and across the medial aspect of the PFC, spanning the medial-orbital (MO), prelimbic (PRL) and anterior cingulate (ACG) areas. All rats alternated between periods of sleep and wake and, while awake, quickly transitioned through behaviors such as locomotion, in-place exploration and grooming. Their breathing patterns varied accordingly, as captured by the nasal pressure signal (Figure 1b, top trace).

Respiratory modulation of brain activity is already apparent upon direct visual inspection of the LFPs at both the OB and PFC (Figure 1b, middle and bottom traces). For reasons that will become apparent as we move forward, inverting the nasal pressure (inhalations up) and OB signals greatly aids in visualizing the alignment between breathing and LFPs and we have adopted this convention throughout the paper. Looking in detail, low-frequency components (< 20 Hz) of the LFP aligned with the nasal breathing cycles can be seen at times of all slow, intermediate and fast breathing (Figure 1c). Crucially, while LFP waves appear consistently aligned with a given phase of the breathing cycle within each rate, this phase does not seem to hold across breathing rates: for slow, intermediate and fast breathing, positive LFP peaks occur by the end of the inhalation phases, halfway through the exhalations, or at the rising phase of the inhalations, respectively. Narrow-band filtering the signals around each breathing frequency highlights the varying phase relationships between brain and respiratory signals across breathing rates (Figure 1c, bottom traces and polar plots).

We set out to characterize the detailed alignment of nasal breathing and frontal brain LFPs and its dependence on breathing rate. We first present results on the low-frequency components of the LFP (Figures 1 and 2) and then on LFP oscillations in the gamma range (Figures 3 and 4). For all the analyses, we will first highlight the results from the cortical electrode most saliently modulated by breathing for each rat (named “PFC”, located in the MO or PRL for 5 and 2 rats respectively; see Methods) and then evaluate the main results across cortical areas.

In our sessions, instantaneous (cycle-by-cycle) breathing rate varied across the 1–11 Hz range, with a bimodality typical of freely-behaving rats [22,79] (Figure 1d). Breathing rate varied with behavior: NREM sleep was characterized by slow breathing (∼1 Hz), grooming by breaths of intermediate rates (∼1.5 to 4 Hz), and exploratory behaviors by faster rates typical of active sniffing (∼6 to 10 Hz) (Figure 1e). The rats modulated this rate rapidly; it took a median time of 0.71 s for the instantaneous rate to change by 2 Hz or more (per-rat medians 0.43–0.97 s).

### Phase difference between frontal brain potentials and nasal breathing is strictly dependent on respiratory rate

Embracing the distinctive rhythmicity of breathing, its synchrony with brain signals is commonly assessed with frequency analysis methods, mainly by computing coherence. In essence, coherence measures the phase difference between two signals at any given frequency and its magnitude gauges how consistent this difference is over the data [58,82]. We developed a method to accurately measure coherence between nasal pressure and LFPs at the breathing rate under conditions of rapidly changing respiratory rates (see Methods, “breathing-brain coherence”). For each rat, we first segmented the data into 1-second windows, sorted them by their peak nasal-pressure frequency into 1-Hz bins (from 1 to 10 Hz; Figure 1f), and then computed the magnitude-squared coherence and phase-difference spectra between nasal pressure and each LFP channel over the (typically hundreds of) windows in each rate bin. From these spectra, we finally extracted the values at the frequency matching the breathing rate, which we named breathing-brain coherence magnitude and phase.

For all breathing rates, and for both the OB and PFC LFPs, the coherence magnitude spectra showed peaks at the exact frequency matching the breathing rate (Figure 1g), evidencing sizable respiratory modulation of frontal brain signals across breathing regimes. Breathing-brain coherence was significant (p < 0.01) for 123 out of 130 combinations of rat, brain region and breathing rate (Table S1). The large coherence of the frontal-brain LFPs to the nasal pressure at all breathing rates allowed us to evaluate the phase difference of this synchrony across the full respiratory repertoire of the rats. Notably, phase difference was dramatically dependent on breathing rate, progressively increasing from ∼0 to 2*π* as the rate increased from 1 to 10 Hz (Figure 1h). This relationship was well captured by a linear regression model (LMM R^2^ = 0.93; slope = 0.194 ± 0.005 π · Hz^−1^; *F*(1,119) = 742.23, *p* < 1e^−6^). Slope was equivalent for OB and PFC, as there was no interaction between brain region and breathing rate (*F*(1,119) = 0.13, *p* = 0.71), and there was a small but significant effect of region on the phase at midpoint rate (PFC − OB = −0.10 ± 0.028 π; *F*(1,119) = 12.60, *p* = 0.0006). Thus, both the OB and PFC LFPs follow the nasal breathing rhythm with a phase difference that is a linear function of the respiratory rate, covering the full 0–2*π* range.

A linear phase-frequency slope is expected when two signals maintain a constant time lag irrespective of frequency [82,83]. In such a case, the value of the slope equals 2 · π · τ, with τ corresponding to the time lag (note that this slope has units of 1/Hz = s). For the slope in our linear model, this would yield *τ* = 96.9 ± 2.4 ms. Independently fitting one linear model for each rat and brain region yielded equivalent predicted time lags for OB and PFC (Figure 1i; OB: 95.0 ± 3.3 ms, PFC: 95.1 ± 9.3 ms; LMM *F*(1,11) = 0.0001, *p* = 0.99). Similar results were observed across PFC areas (Figure S1a–c). Overall, the analyses above suggest that the respiratory components of the frontal LFPs follow nasal pressure with a constant delay of approximately 90 to 100 ms.

### Frontal brain potentials align with nasal breathing with constant latency across all respiratory rates

To directly quantify the alignment of the LFP and breathing in the time domain, we turned to cross-correlation analysis, which measures the linear correlation between two signals as a function of their time lag [58]. We obtained the cross-correlograms between the LFPs and nasal pressure for the same 1-second windows of data sorted by breathing rate as before.

The analysis revealed prominent temporal alignment of both OB and PFC LFPs to the nasal-pressure signal for all breathing rates (Figure 2a). Maximum correlation R was significant (p < 0.01) for all 130 combinations of rat, brain region and breathing rate (Table S1). The shape of the cross-correlation functions varied with breathing rate, progressing from wider to narrower peaks as the rates grew from 1 to 10 Hz. Strikingly, they showed a distinct positive peak at LFPs lagging nasal pressure by ∼100 ms for all breathing rates (Figure 2a, vertical dashed lines). Indeed, the lag of the correlation peaks appeared constant when plotted against respiratory rate for both OB and PFC (Figure 2b). A linear regression model revealed no effect of breathing rate on peak lag (LMM R^2^ = 0.13; slope = −1.15 ± 0.75 ms · Hz^−1^; *F*(1,126) = 1.07, *p* = 0.30), and no interaction between rate and brain region (*F*(1,126) < 0.01, *p* = 0.99). The overall model estimate for lag at midpoint rate was 100.1 ± 4.6 ms, and was significantly smaller for PFC (PFC − OB = −10.8 ± 4.4 ms; *F*(1,126) = 5.97, *p* = 0.016). Thus, slow LFP components in both the OB and PFC follow nasal pressure with constant time lags across the whole range of respiratory rates. The mean lags over rates closely match the mean delay predicted by the coherence phase-frequency slope above (Figure 2c; OB: 103.3 ± 2.3 ms, PFC: 94.7 ± 6.9 ms; no significant difference between brain regions, LMM *F*(1,11) = 2.0, *p* = 0.19), confirming that the strong dependence of phase on breathing rate is explained by a fixed time lag between frontal-brain LFPs and nasal pressure. Similar results were observed across PFC areas (Figure S1d–f).

Fixed latencies are typical of brain activity evoked by sensory stimuli [64,65,84]. We thus wondered whether studying the breathing-brain signals as chained evoked potentials from respiratory events could aid in characterizing the modulation of frontal brain activity by breathing. For this, we first segmented the nasal pressure signal into individual respiratory cycles and sorted them into 1-Hz bins according to their instantaneous rate (Figure 2d). We then used the times of inhalation peaks as events to obtain event-triggered averages of both the nasal pressure and LFP time series (hereon called “inhalation-triggered averages” or “ITAs”). The average shape of the breathing cycle waveforms changed with instantaneous rate, as revealed by the nasal-pressure ITAs, from asymmetric with prominent inhalations but slow and shallow exhalations at low rates to more symmetrical sinusoidal waves at higher rates (Figure 2e). As expected, the latency from inhalations to subsequent peak exhalations decreased with increasing breathing rate as breathing cycles got shorter (Figure 2f, g, top). Approximating the rate-latency curve with a linear fit confirmed a significant effect of breathing-cycle rate on inhalation-to-exhalation latency (LMM R^2^ = 0.72; slope = −33.8 ± 2.5 ms · Hz^−1^; *F*(1,68) = 177.92, *p* < 1e^−6^), with pronounced slopes across rats (Figure 2i; slope different from 0, *t*(6) = −11.3, *p* = 0.00003).

The ITAs of both OB and PFC LFPs revealed large and consistent potentials aligned to inhalation times across all breathing-cycle rates (Figure 2f, middle and bottom; RMS at 0– 500 ms latency was significant at p < 0.01 for 129 out of 130 combinations of rat, brain region and breathing rate; Table S1). A first prominent positive peak (“P1”) appeared at a latency of ∼100 ms regardless of rate. A negative peak (“N1”) ensued but, in contrast with P1, its latency appeared to depend on breathing-cycle rate, steadily moving closer in time to P1 as the rate increased (Figure 2g, middle and bottom). Fitting a linear model to the P1 latencies confirmed no significant effect of breathing-cycle rate (LMM R^2^ = 0.35; latency at midpoint rate = 109.3 ± 4.6 ms; slope = −1.02 ± 0.41 ms · Hz^−1^; *F*(1,126) = 1.19, *p* = 0.28), brain region (PFC – OB = −2.98 ± 2.42 ms; *F*(1,126) = 1.51, *p* = 0.22) or their interaction (PFC − OB = −0.73 ± 0.82 ms · Hz^−1^; *F*(1,126) = 0.80, *p* = 0.37). In contrast, there was a strong effect of breathing-cycle rate on N1 latencies (LMM R^2^ = 0.80; slope = −30.4 ± 1.3 ms · Hz^−1^; *F*(1,126) = 288.51, *p* < 1e^−6^), with effects for brain region on latency at midpoint rate (PFC − OB = −18.5 ± 7.7 ms; *F*(1,126) = 5.80, *p* = 0.017) and interaction between rate and region (PFC − OB = 5.92 ± 2.68 ms · Hz^−1^; *F*(1,126) = 4.89, *p* = 0.029).

The similarity between the rate-latency curves for LFP N1 peaks with the one for exhalation peaks (compare N1 to Exh in Figure 2g) suggested that the negative phases of the evoked potentials could be related to exhalations. Indeed, subtracting the latencies of exhalations from those of LFP peaks revealed that N1 peaks had a constant latency to mean exhalation-peak times across breathing-cycle rates (Figure 2h; LMM R^2^ = 0.049; latency at midpoint rate = 94.3 ± 5.3 ms; slope = 3.59 ± 1.83 ms · Hz^−1^; *F*(1,126) = 0.090, *p* = 0.76), with no effect of brain region on latency at midpoint rate (PFC − OB = −18.2 ± 10.5 ms; *F*(1,126) = 3.00, *p* = 0.09) or interaction (PFC − OB = 5.56 ± 3.67 ms · Hz^−1^; *F*(1,126) = 2.30, *p* = 0.13). In contrast, re-aligning the P1 peaks to exhalation times results in latencies strongly dependent on breathing rate (LMM R^2^ = 0.68; slope = 33.0 ± 2.0 ms · Hz^−1^; *F*(1,126) = 132.17, *p* < 1e^−6^), with no effects for brain region on latency at midpoint rate (PFC − OB = 0.52 ± 11.4 ms; *F*(1,126) = 0.0021, *p* = 0.96) or interaction (PFC − OB = −1.09 ± 3.97 ms · Hz^−1^; *F*(1,126) = 0.075, *p* = 0.78).

To directly compare the alignments of P1 and N1 peaks to inhalations and exhalations, we computed the rate-latency slope for each combination of rat, peak identity (P1 and N1) and respiratory time-reference (inhalations and exhalations) (Figure 2i). For both OB and PFC, the slopes of P1 latency vs. rate were not different from 0 across rats when measured from inhalations (one-sample t-tests; OB: *t*(5) = −1.74, *p* = 0.14; PFC: *t*(6) = −0.83, *p* = 0.43). Instead, slopes were pronouncedly positive if measuring P1 latency to exhalations (OB: *t*(5) = 9.11, *p* = 0.0003; PFC: *t*(6) = 9.48, *p* = 0.00008). Slopes were indeed larger when aligning P1 to exhalations than to inhalations (paired t-tests; OB: *t*(5) = 9.72, *p* = 0.0002; PFC: *t*(6) = 11.30, *p* = 0.00003). Conversely, N1 rate-latency slopes were not different from 0 when measured from exhalations (OB: *t*(5) = 0.28, *p* = 0.79; PFC: *t*(6) = 1.55, *p* = 0.17), but were pronouncedly negative when referencing to inhalations (OB: *t*(5) = −17.67, *p* = 0.00001; PFC: *t*(6) = −15.84, *p* = 0.000004). Slopes were indeed steeper (more negative) when aligning N1 to inhalations than to exhalations (paired t-tests; OB: *t*(5) = 9.72, *p* = 0.0002; PFC: *t*(6) = 11.30, *p* = 0.00003). Thus, P1 LFP peaks show constant latency to inhalations independent of breathing rate, while N1 LFP peaks do so to exhalations. Directly contrasting the mean latencies of P1 to inhalations with those of N1 to exhalations reveals no significant differences for either OB and PFC (Figure 2j; paired t-tests; OB: *t*(5) = 0.66, *p* = 0.54; PFC: *t*(6) = 2.11, *p* = 0.08). Thus, nasal inhalations and exhalations respectively evoke positive and negative slow potentials throughout the frontal brain, with equivalent constant latencies of ∼100 ms independently of breathing rate.

Leveraging what we have learned so far to realign the signals reveals an astounding match between the average nasal-pressure and LFP waveforms across all breathing-cycle rates (Figure 2k, see Figure S2 for all rats). This superimposition was achieved by (1) inverting the original nasal pressure signal, thus leaving inhalations positive, (2) inverting our superficial-OB LFP, (3) z-scoring each average waveform and — crucially — (4) delaying the nasal pressure signal by 100 ms. Thus, the slow frontal brain LFP components aligned to nasal breathing behave on average as scaled, time-delayed copies of the nasal-pressure waveform.

Feedforward propagation of activity from the OB to the PFC would be expected to add an extra latency of ∼20 ms in rats [52], yet we found no consistent delay between PFC and our superficial OB signal. LFPs can reflect activity originating at a distance from the recording electrode, as potentials from large synchronous neuronal populations can propagate via volume conduction [85–87]. While our bulbar and cortical potentials cannot be directly related through volume conduction since they are of opposite polarity, early respiratory components of the OB LFP themselves are positive in deeper bulbar layers [88–90]. To verify that our data reflects true respiratory modulation of cortical activity and not volume conduction from the deep OB, we obtained differential LFPs (∇LFP, current flow density) — which effectively removes common distal potentials [87,91,92] — from neighboring electrode pairs within the PFC. Cross-correlation and ITA analyses of PFC ∇LFP thoroughly reproduced our findings above for the LFP, confirming that the breathing cycle modulates local currents in the PFC with a fixed latency of ∼100 ms largely independent of breathing-cycle rate (Figure S3 and Table S2). The comparable latencies we observed in the OB may be a consequence of breathing-locked potentials in its superficial layers being themselves delayed from a true feedforward wave detected deeper in the bulb [88–90] (see Discussion).

### Nasal inhalations and exhalations evoke bursts of fast and slow gamma oscillations, respectively

LFP oscillations in the gamma frequency range (∼40–100 Hz) are prominent in both the OB and PFC, and their power has been shown to be modulated by the breathing rhythm [7,11,12,16,19,37,56,93–96]. We set out to characterize the detailed alignment of local gamma oscillations to the nasal breathing cycle across the full range of rat respiratory rates. High frequency activity was readily apparent on the OB and PFC LFPs of our rats, riding over the slower respiratory modulation of the brain potentials (Figure 3a, b, top). During slow breathing (Figure 3a, ∼2–3 Hz), faster gamma bursts (∼60–100 Hz) were commonly observed to coincide with the inhalation-locked LFP peaks (“P1” above), while slower gamma activity (∼40–60 Hz) could be seen riding the LFP valleys associated with the exhalation phases (“N1” above). The power of these fast-and slow-gamma bursts appeared variable across breathing cycles, with some cycles showing clear bursts of both types and others only one of them or none (see spectrograms in Figure 3a). During sniffing (Figure 3b, ∼7–8 Hz), only fast-gamma bursts were typically observed and, as for slow breathing, they aligned with the inhalation-locked LFP peaks. Overall, modulation of the broad gamma envelope (50–100 Hz) was significant (p < 0.01) for 108 out of 130 combinations of rat, brain region, and breathing rate.

We next performed ITAs of the OB and PFC gamma spectrograms across breathing-cycle rates. The average power spectral density aligned to the nasal inhalations revealed distinct fast-gamma bursts coinciding with the P1 peak of the LFP (∼100–150 ms from inhalation peaks) for OB and PFC for both slow and fast breathing (2- and 7-Hz breathing-cycle rate bins, Figure 3c, d; see Figure S4 for all rats). The instantaneous frequency of the average fast-gamma bursts slows down during their short duration, from ∼100 to ∼60 Hz in under 50 ms. Slow-gamma bursts could be observed at longer latencies, closer to that of the N1 LFP peak, but only for the slow breathing cycles (Figure 3c). Intriguingly, while respiratory fast-gamma bursts were observed in the OB and PFC of all rats, slow-gamma bursts were absent or too weak in both brain regions for two of them (Figure S4; rats excluded from all further analyses of slow gamma). Looking in detail across breathing-cycle rates, this respiratory slow gamma was most salient for cycles in the ∼2–4 Hz range (Figure S5a). Furthermore, the peak frequency of the fast-gamma burst was lower in PFC vs. OB, most noticeably for fast breathing-cycle rates (Figure S5b).

To evaluate in detail the alignment of these gamma activities to nasal breathing, we filtered the LFPs at narrow frequency bands centered at the observed fast (70–90 Hz) and slow (50– 55 Hz) gamma ranges. We then obtained their envelopes, capturing the instantaneous amplitude of the band-filtered signals, and computed their ITAs. Across the whole 1–10 Hz range of breathing-cycle rates, fast-gamma amplitude peaked shortly after 100 ms from inhalation peaks for both OB and PFC (Figure 3e). A linear model revealed that the latency from inhalations of fast-gamma envelope peaks was only marginally dependent on breathing-cycle rate (Figure 3f; LMM R^2^ = 0.15; slope = −0.89 ± 0.45 ms · Hz^−1^; *F*(1,126) = 3.92, *p* = 0.050; latency at midpoint rate = 115.4 ± 0.9 ms), with no effect of brain region on latency at midpoint rate (PFC − OB = 2.02 ± 1.76 ms; *F*(1,126) = 1.31, *p* = 0.25) or interaction (PFC − OB = −1.00 ± 0.61 ms · Hz^−1^; *F*(1,126) = 2.64, *p* = 0.11). Modulation of the fast-gamma envelope at 0– 200 ms from inhalation was significant (p < 0.01) for 115 out of 130 combinations of rat, brain region, and breathing rate (Table S1).

As mentioned above, respiratory slow-gamma bursts are detectable over a narrower range of slow breathing-cycle rates. Looking in detail within this range (2–4 Hz), slow-gamma envelope peaked progressively later in time from inhalations as the breaths got longer (Figure 3g). The latency from inhalations of slow-gamma envelope peaks was indeed strongly dependent on breathing-cycle rate (Figure 3h; LMM R^2^ = 0.95; slope = −79.8 ± 4.3 ms · Hz^−1^; *F*(1,40) = 351.79, *p* < 1e^−6^; n = 4 and 5 rats for OB and PFC), with no effect of brain region on latency at midpoint rate (PFC − OB = −3.69 ± 4.26 ms; *F*(1,40) = 0.75, *p* = 0.39) or interaction (PFC − OB = 2.58 ± 5.83 ms · Hz^−1^; *F*(1,40) = 0.20, *p* = 0.66). As before for the N1 LFP peak, this pattern suggested that slow-gamma bursts could be aligned to exhalations instead of inhalations. Obtaining their latencies from the times of exhalation peaks revealed that slow-gamma bursts are indeed time-locked to nasal exhalations with a constant latency of ∼110 ms (Figure 3h; LMM R^2^ = 0.29; latency at midpoint rate = 113.5 ± 11.2 ms; slope = −11.6 ± 8.8 ms · Hz^−1^; *F*(1,40) = 1.72, *p* = 0.20), with no effect of brain region on latency at midpoint rate (PFC − OB = −2.75 ± 8.87 ms; *F*(1,40) = 0.096, *p* = 0.76) or interaction (PFC – OB = 10.5 ± 12.1 ms · Hz^−1^; *F*(1,40) = 0.75, *p* = 0.39). Modulation of the slow-gamma envelope at 0– 200 ms from exhalation was significant (p < 0.01) for 44 out of 45 combinations of rat, brain region, and breathing rate (Table S1).

To verify that gamma oscillations of local PFC currents are time-locked to nasal airflow, we repeated the analyses above in ∇LFP signals (Figure S6). Both the fast- and slow-gamma bands of PFC current flow were robustly modulated by the breathing cycle (significant modulations more prevalent in MO than in PRL or ACG electrode pairs; Table S2). ITA analysis of the gamma ∇LFP thoroughly reproduced the results obtained for the LFP, confirming that inhalations and exhalations respectively evoke bursts of fast and slow gamma activity in local PFC currents with a fixed latency of ∼100–150 ms that is largely independent of breathing-cycle rate.

### Respiratory gamma bursts in the PFC are not directly propagated from the OB

We wondered whether the respiratory gamma bursts in the PFC are directly propagated from gamma oscillations in the OB, or if they reflect locally generated oscillations timed by the breathing cycle. Close observation of the OB and PFC signals revealed that the relative strength of their respiratory gamma bursts could vary substantially across breathing cycles (Figure 4a; see also Figure 3a, b), hinting at some independence in their generation. The saliency and evoked-response behavior of the gamma bursts allow for estimating their amplitude at each breathing cycle in time windows locked to inhalations and exhalations (Figure 4a and Methods). We took advantage of this to quantify the cycle-by-cycle amplitude correlation between OB and PFC of γ_fast_ and γ_slow_ for slow breathing (2–4 Hz) and of γ_fast_ for sniffing (6–8 Hz; Figure 4b), keeping only cycles for which salient (> 2 z) respiratory gamma was detected in the OB. PFC gamma-burst amplitudes were significantly but only moderately correlated with their OB counterparts, with OB amplitude explaining about 10% of the variance in PFC amplitude overall (Figure 4b, d; R^2^, mean ± sem (n rats): 2–4 Hz, γ_fast_ 0.09 ± 0.03 (6), γ_slow_ 0.13 ± 0.04 (4); 6–8 Hz, γ_fast_ 0.04 ± 0.02 (6); all p ≤ 0.018). To put these values in context, we contrasted them with correlations between pairs of cortical electrodes, for which the LFP signals can be expected to more closely match each other through short-distance circuit transmission or common current sources. Respiratory-gamma amplitudes were robustly correlated for PFC electrode pairs, with one typically explaining 70% of the variance of the other (Figure 4c, d; PFC > 2 z; R^2^: 2–4 Hz, γ_fast_ 0.68 ± 0.04 (5), γ_slow_ 0.77 ± 0.06 (4); 6–8 Hz, γ_fast_ 0.56 ± 0.08 (5); all p < 1e^−6^). R^2^ values were significantly smaller for OB-PFC vs. PFC-PFC’ pairs (paired t-tests: 2–4 Hz, γ_fast_ *t*(4) = 7.94, *p* = 0.0014, γ_slow_ *t*(3) = 10.73, *p* = 0.0017; 6–8 Hz, γ_fast_ *t*(4) = 5.59, *p* = 0.0050).

The results above suggest that respiratory gamma bursts in the PFC are not a direct copy of those in the OB. Either they are locally generated or, if propagated from the OB, the gain of this transmission is dramatically variable across breathing cycles. An independent signature of direct wave propagation would be strict phase coupling between the PFC and OB gamma waves. Visually, this phase relationship appeared variable across breathing cycles (Figure 4a, insets). We measured OB-PFC magnitude-squared coherence of the gamma bands in the same time windows above (Figure 4e). PFC respiratory gamma bursts were significantly but moderately coherent with their OB counterparts, with the OB fast-and slow-gamma bands explaining about 10 and 20% of the variance in the corresponding PFC signals (Figure 4e; MSC, mean ± sem (n rats): 2–4 Hz, γ_fast_ 0.07 ± 0.03 (6), γ_slow_ 0.23 ± 0.06 (4); 6–8 Hz, γ_fast_ 0.07 ± 0.04 (6); all p ≤ 0.001). These values were markedly lower than those between PFC electrode pairs (Figure 4e; MSC: 2–4 Hz, γ_fast_ 0.91 ± 0.02 (5), γ_slow_ 0.92 ± 0.03 (4); 6–8 Hz, γ_fast_ 0.88 ± 0.04 (5); all p ≤ 0.001, cycle-shuffling surrogates; paired t-tests: 2–4 Hz, γ_fast_ *t*(4) = 20.91, *p* = 0.000031, γ_slow_ *t*(3) = 12.42, *p* = 0.0011; 6–8 Hz, γ_fast_ *t*(4) = 16.12, *p* = 0.000087). Overall, our results suggest that breathing cycles coordinate the local emergence of fast and slow gamma waves in PFC circuits, with constant latency to nasal inhalations and exhalations. Interestingly, these bursts are neither directly imposed by their OB counterparts nor completely independent from them, as revealed by the moderate but significant phase coherence and co-modulation of their amplitudes.

## Discussion

In this work, we demonstrate that respiratory components of the LFP across the OB and PFC (MO, PRL and ACG) of the rat brain align with constant latencies of ∼100 ms to the cyclic fluctuations in intranasal pressure/airflow resulting from natural breathing patterns. These breathing-brain signals thus behave as is expected for waves of evoked sensory activity. We further unveil distinct activity patterns evoked by ortho- and retronasal airflow: LFP peaks and fast-gamma bursts time-locked to inhalations and LFP troughs and slow-gamma bursts aligned to exhalations. Our results offer new perspectives on the mechanisms and functions of respiratory modulations of brain activity and propose methodological avenues for their quantitative analysis.

Current evidence supports that the respiratory modulations of LFPs in the frontal rodent brain are largely dependent on activation of the olfactory system [6,7,16,38,40,41]. OSNs are distributed along the convoluted ethmoturbinates supporting the OE [39,97,98], and are sensitive to both pressure and airborne odorants [45,46]. The transformation of nasal airflow into OSN and OB-glomeruli activation patterns results from the complex interplay of flow dynamics, chemical composition, nasal anatomy, and topology of olfactory receptor expression [47,81,99–102]. Rodent studies have reported ∼50–150 ms mean delays from inhalation onsets to OE activations [11,44,99], suggesting that the bulk of the OB/PFC latencies we observed may happen already at the level of sensory transduction. These long delays likely result from a combination of slow/nonuniform airflow reaching the OE in the ethmoturbinates with the slow nature of the biochemical signaling triggered by the G-protein-coupled olfactory receptor (OR) activation by both pressure and odorants in the OSNs [45,46]. Similar delays were reported in the OB, with some studies already showing largely constant latencies of LFP and single units to inhalations across nasal-breathing rates [60–62,99,100].

We detected two families of LFP components evoked by nasal airflow across the OB and PFC: slow potentials matching the pressure waveforms in the nose (Figures 1 and 2) and fast oscillation bursts time-locked to inhalations and exhalations (Figure 3 and 4). Extracellular field potentials at the analyzed frequency range (< 100 Hz) result mainly from the summation of synaptic currents in correlated populations of neurons [85,103]. The slow evoked potentials at the OB and PFC likely sum contributions from feedforward synaptic inputs propagated from the OE with feedback stemming from local neuronal activations.

The fact that evoked potentials from inhalations and exhalations are of opposite polarity suggests that breathing phases push the local networks into fundamentally different alternating states. Interestingly, already at the OE, field potentials reverse polarity from inhalation to exhalation [44] and OB glomeruli and projection neurons have preferred activation phases that tile the whole respiratory cycle [21,47,99,100,104,105]. It thus seems that the brain responds to nasal breathing not with pulsating activation from inhalations only, but rather by switching between distinct activity patterns associated with ortho- and retronasal airflow. This picture is strengthened by our observation that inhalations and exhalations trigger different patterns of local network activity in the form of fast and slow gamma oscillation bursts appearing shortly (within 50 ms) after the respective evoked potential peaks. Respiratory modulation of gamma oscillations has been detected across the mammalian brain [7,12,16,19,23,37,56,95,96]; distinct fast and slow respiratory gamma activities have been described in the rodent OB and olfactory and orbitofrontal (OFC) cortices, and are thought to arise from different circuit mechanisms [93,94]. We found cases of robust modulation of each gamma band in MO, PRL and ACG, in both LFP and ∇LFP. However, there was a clear regional pattern, with salient and significant respiratory gamma more prevalent in MO current flows (Table S2). Neither the precise time-locking of gamma bursts to nasal airflow nor the emergence of slow respiratory gamma in PFC circuits has, to the best of our knowledge, been documented before.

Our results posit respiratory signals in the frontal brain of rodents as evoked sensory potentials, which trigger local patterns of network activity along their way. What functional roles could they subserve? To begin with, they provide distributed brain circuits with a salient reference to the ongoing breathing pattern, directly linked to nasal airflow fluctuations. While this signal is delayed by ∼100 ms, it is normal for brain circuits to have no direct access to external references, and use local patterns of population activity for latency/phase coding instead [64,106]. Neurons across the brain could thus encode information in their phase of firing with respect to the local — delayed — copy of the respiratory rhythm. This distributed reference signal could contribute to sensory coding, not only for olfactory stimuli [20,21,47,107,108], but also for binding across sensory modalities [109]. Binding could be particularly relevant in rodents, which synchronize a broad set of sensorimotor patterns with the respiratory rhythm during active behaviors [79,110–113]. The alternating network states triggered by inhalations and exhalations could play a fundamental role in the differential processing of ortho- vs. retronasal olfactory information in the brain [114–118]. At the OB, odorants recruit qualitatively similar glomerular maps independent of airflow direction [119–121], but the detailed respiratory timing of circuit activity helps disentangle flow route from odor identity and concentration [20,47,105,107]. Of note, while we report on mean activations, there could be valuable information encoded in the cycle-by-cycle variations of evoked activity patterns (such as that observed for gamma bursts, Figure 3a, b and Figure 4). As we have discussed before [5], the functions of global respiratory oscillations in brain activity could exceed sensory coding, through evolutionary exaptation of this perennial rhythm [122].

A variety of techniques are used to monitor the breathing pattern in rodents, including nasal pressure/airflow [28,34,80], whole body plethysmography [123], nasal air temperature [38,44,61], respiratory muscle activity [62], chest belts [36,46,124] and populational OE or OB activity [11,16,93,125]. We measured air pressure in the dorsal meatus of the nasal cavity, a main passage for both inhalation and exhalation airflow [102]. In the laminar regime of the rat nose, pressure and airflow are linearly related, such that our breathing signal can be interchangeably interpreted as instantaneous local pressure or airflow [81,126]. Its high sensitivity and fidelity make intranasal pressure a privileged signal for detailed analyses of the breathing pattern [21,79,80,127–129]. We do not expect our results to immediately apply to recordings with all other breathing measures, as any phase or time differences between respiratory signals should alter their alignment with brain activity. Extra caution should be exercised when recording nasal air temperature with thermocouples or thermistors, as the phase lags imposed by their low-pass filtering of the nasal air temperature can result in time lags and signal attenuations that are a non-straightforward function of breathing rate and airflow speed [11,80,130–132]. The electroolfactogram (EOG; measuring OE populational activity) stands out as an interesting complementary reference for breathing-brain signals, as it bypasses the latency from sensory transduction and should be stable, not being the subject of top-down modulations [11,16]. Simultaneously quantifying the alignment of brain activity to all these breathing signals in the same animals would be invaluable to reconcile past and future results across the literature.

Acknowledging time-locking between neural and breathing signals can guide the choice of analysis methods. For studying possible modulations of their alignment across conditions, measures of phase difference should be avoided when breathing rate is not constant, as spurious phase shifts can be expected for even modest changes in rate (see Figure 1h).

Synchrony metrics in the time domain (latency to respiratory events, cross-correlation lags) would be recommended instead. We note that assessing the phase-locking of neuronal spiking to a local-circuit breathing reference such as LFP or neuronal population activity is a valid and most interesting avenue to pursue, as the local reference would already absorb the delay from nasal airflow. As for comparing the magnitude of breathing-evoked potentials across conditions, phase-based techniques — such as magnitude-squared coherence or phase locking value — give only indirect estimates, as they measure synchrony to the breathing signal relative to other components at the same frequency. This is especially critical when breathing rate is not constant within the analysis window, as phase-locking would be smeared across the frequency axis [58,83]. Our rate-resolved coherence method (Figure 1g) helps minimize this latter confound by measuring coherence over short windows of the same breathing rate. Event-triggered averages (event-related potentials) are recommended instead, as they give unbiased estimates of the true average waveform, whose variability decreases with the number of events included [58,84]. ITA allows for pooling breathing cycles by conditions (behavioral, physiological, etc.), even of varying rate, and contrasting the amplitude of neural signals (LFP, power of high-frequency oscillations, spike rate) at the observed fixed latency (i.e. 100 ms). We note that most of the methods in this work do not allow for instantaneous measurement of modulation strength, since they require averaging over a number of time windows/breathing cycles. Respiratory gamma bursts can be salient enough from baseline for their amplitude to be estimated cycle by cycle (Figure 4), but we note that this measure is noisier than cycle averages because of variability in their latency and background activity. Alternative methods are worth exploring, such as matching the complete breathing cycle waveform (time-delayed as needed) to that of the neural signal [14,133] or estimating response functions from continuous or event-based regression [134,135].

Analyzing brain signals associated with natural breathing patterns under a stimulus-response framework poses some challenges. Airflow in the nose is a continuous stimulus waveform, with no obvious onsets. We chose the peak inhalation airflow as a salient landmark that can be unambiguously detected for the vast majority of cycles across all rates. Exhalation peaks are less defined for individual slow cycles, so that they were extracted from the average pressure waveforms when aligning brain signals to them. Alternative references can be explored, such as transition times between inhalations and exhalations.

Furthermore, because of the uninterrupted cyclical nature of the stimulus, there are no true baselines available for analysis, and response waveforms from adjacent cycles can overlap in time. Also, most of our results are from averages over hundreds of cycles or time windows, such that they do not explore potentially interesting variability in the responses within breathing-rate bins.

Feedforward propagation of activity should impose additional delay at each synapse. Direct stimulation of the OB or its projection axons evokes short-latency responses in the piriform (< 10 ms [136]), entorhinal (< 20 ms [137]), and prefrontal (∼20 ms [52]) cortices, yet we found no consistent differences in latency between OB and PFC. Several results demonstrate that this is not due to PFC LFPs being volume conducted from the OB: (1) Cross-correlation and ITA analysis of PFC ∇LFP signals revealed salient local currents time-locked to nasal airflow with fixed ∼100 ms latencies across breathing rates (Figure S3); (2) Prominent bursts of ∇LFP fast and slow gamma were likewise locked to inhalations and exhalations with equivalent latencies (Figure S6); (3) Fast-gamma bursts differed in frequency content between OB and PFC LFPs (Figure S5b) and (4) Neither the amplitude nor the phase of the fast-and slow-gamma bursts in the PFC strictly match their OB counterparts cycle-by-cycle (Figure 4). Close examination of laminar profiles reported for the respiratory component of the rodent LFP reveals that while deep layers (below the mitral cell layer) exhibit an early positive component following inhalations, more superficial potentials are dominated by a negative potential that peaks visibly later in the breathing cycle (see Figure 1B in [88] and Figure 7B and S5F in [89] for rats and Figure S2B in [90] for neonatal mice). This late component could be driven by propagation within the OB circuit or feedback activity from olfactory cortices [138–140] but its exact nature has not been established to the best of our knowledge. The added delay from deep OB layers to our superficial recording site could explain why OB and PFC latencies are comparable in our LFP data. We expect breathing-evoked potentials in the PFC to be delayed with respect to the early deep wave of OB activation, compatible with feedforward circuit transmission. Further studies will be needed to map the precise latencies of respiratory evoked signals across OB and PFC layers and PFC areas through analyses of current source density or local neuronal spiking [85,91]. We further observed equivalent latencies at the OB and PFC for both fast- and slow-gamma bursts, with partial but significant phase coherence between them. Long-range synchrony between respiratory-driven gamma in the OB, thalamus and frontal cortices has been shown before [12,37,94,141,142], suggesting that circuit mechanisms could contribute to binding their emergence in time.

We limited our recordings to the dark phase of the light cycle, when rats are more active. Accordingly, the rats were predominantly awake and active during the sessions. Nevertheless, all rats interleaved long periods of NREM sleep (18% of valid recording time on average, range 10.6–21.7%). Putative REM episodes were few and short-lived, as expected during the active period [143]. As such, our 1-Hz breathing rate bins mostly represent NREM-sleep breathing cycles (see Figure 1e). Extending sessions to the light phase would be interesting to contrast against breathing-brain signals during REM sleep, during which rat breathing patterns are known to differ from NREM [127]. Our study used only male subjects, a limitation due to animal availability, such that our results cannot be directly extrapolated to females. Some studies have contrasted rodent breathing patterns between males and females and reported no salient effects of sex or sex hormones [113,128,144,145] (but female mice show more sleep-disordered breathing [146]). While most studies of breathing-brain signals are single sex or do not inform on sex differences, those that compare the sexes report equivalent results for males and females [10,35,75,147]. In the rat OB, male and female rats exhibit equivalent gamma oscillations during free behavior, but an increase in gamma power during odor presentation is weaker in females [148], while in anesthetized mice, odors activate more OB glomeruli and faster in females [149]. Overall, we do not expect our main findings to be qualitatively different in females, but quantitative differences in the amplitude and exact latency of the breathing-evoked signals could exist, particularly during exposure to strong odors. More studies with balanced sex design are needed to unveil possible sex effects in the respiratory modulation of cortical circuits.

Our results further support that breathing does not act by entraining endogenous oscillations in the rodent brain, as we advanced in [5]. We show that PFC LFPs exhibit robust synchrony to nasal airflow across the full 1–10 Hz range of rat breathing rates. Because phase-locking between weakly-coupled oscillators can only emerge if their natural frequencies are close [69,150] and no single known neuronal oscillation spans this range [66], breathing would need to ‘jump’ between recruiting different ones such as delta (1–4 Hz) and theta (6–10 Hz), depending on its rate. However, both delta and theta have been shown to independently coexist with breathing-brain rhythms [34,36,56,57]. This unlikely scenario is made even less plausible by our demonstration that the phase difference between breathing and brain signals changes linearly with rate, spanning the full 0–2π (Figure 1h). Coupled oscillators should exhibit a narrow range of preferred phase differences across their frequency band of entrainment instead [69,73,74,150]. This result is thus incompatible with the entrainment hypothesis but fully expected for evoked responses with fixed latencies [72–74,82,83]. Whether brain signals synchronous to periodic sensory inputs reflect sequences of evoked responses or entrainment of endogenous cortical oscillations is extensively debated in other systems, particularly in the human neurophysiology literature [71,73,74,151–155].

While our results are fully in line with the evoked scenario for neuronal populations in the rat PFC, we do not rule out the possibility that breathing-brain signals at other scales (e.g. unit activity, global brain networks), brain regions (e.g. subcortical) or species (e.g. humans) could reflect true entrainment of local oscillations.

## Conclusions

Our results support that population activity in the rat PFC is modulated by synaptic activity originated by cyclic stimulation of the olfactory epithelium in the nose. A respiratory cortical rhythm would thus emerge from the chaining of sensory evoked potentials with fixed latency to the nasal airflow oscillation. This signal lies at a fascinating crossroads between interoception and exteroception, following the vital internal rhythm of breathing while adding relevant external information on its reafferent route through the nose. Endogenous neuronal oscillations have been linked to a vast array of functions. Our proposal that breathing does not act by recruiting them but rather by imposing its own pattern can guide future work aimed at unveiling unique functional roles for this salient modulation of cortical circuits.

## Acknowledgments

This work was supported by the Brazilian National Council for Scientific and Technological Development (CNPq 461735/2014-8, 305445/2022-7, 444084/2024-0, 446443/2025-5), the Brazilian Coordination for the Improvement of Higher Education Personnel (CAPES 88887.895115/2023-00), the Serrapilheira Institute (Public call N°1-2018, Brazil), INCT-NeuroComp (CNPq 408389/2024-9), the Advanced Knowledge Center in Immersive Technologies (AKCIT PPI IoT MCTI EMBRAPII 057/2023), Fundação de Amparo à Pesquisa do Estado de Goiás (FAPEG 64448878/2024) and the Alexander von Humboldt Foundation. We thank Davi Drieskens for his contribution in collecting the data.

