## Supplementary Figures and Tables for "Nasal inhalations and exhalations evoke distinct prefrontal-cortex responses with constant latency"

* Equal contribution

### Supplementary material

#### Supplementary figures

Figure S1. Alignment of LFPs to nasal breathing across PFC areas.

Figure S2. Average LFP waveforms aligned to inhalations across rats and breathing rates.

Figure S3. Time-locking of cortical current flow (∇LFP) to the nasal breathing cycle.

Figure S4. Average gamma spectrogram aligned to inhalations across rats.

Figure S5. Respiratory fast- and slow-gamma bursts across breathing-cycle rates.

Figure S6. Time-locking of fast- and slow-gamma bursts of cortical current flow (∇LFP) to the nasal breathing cycle.

#### Supplementary tables

Table S1. Statistical significance of metrics per subject, brain region and breathing rate.

Table S2. Statistical significance of metrics for cortical current flow (∇LFP) per subject, brain region and breathing rate.

#### Supplementary Methods

Current flow density.

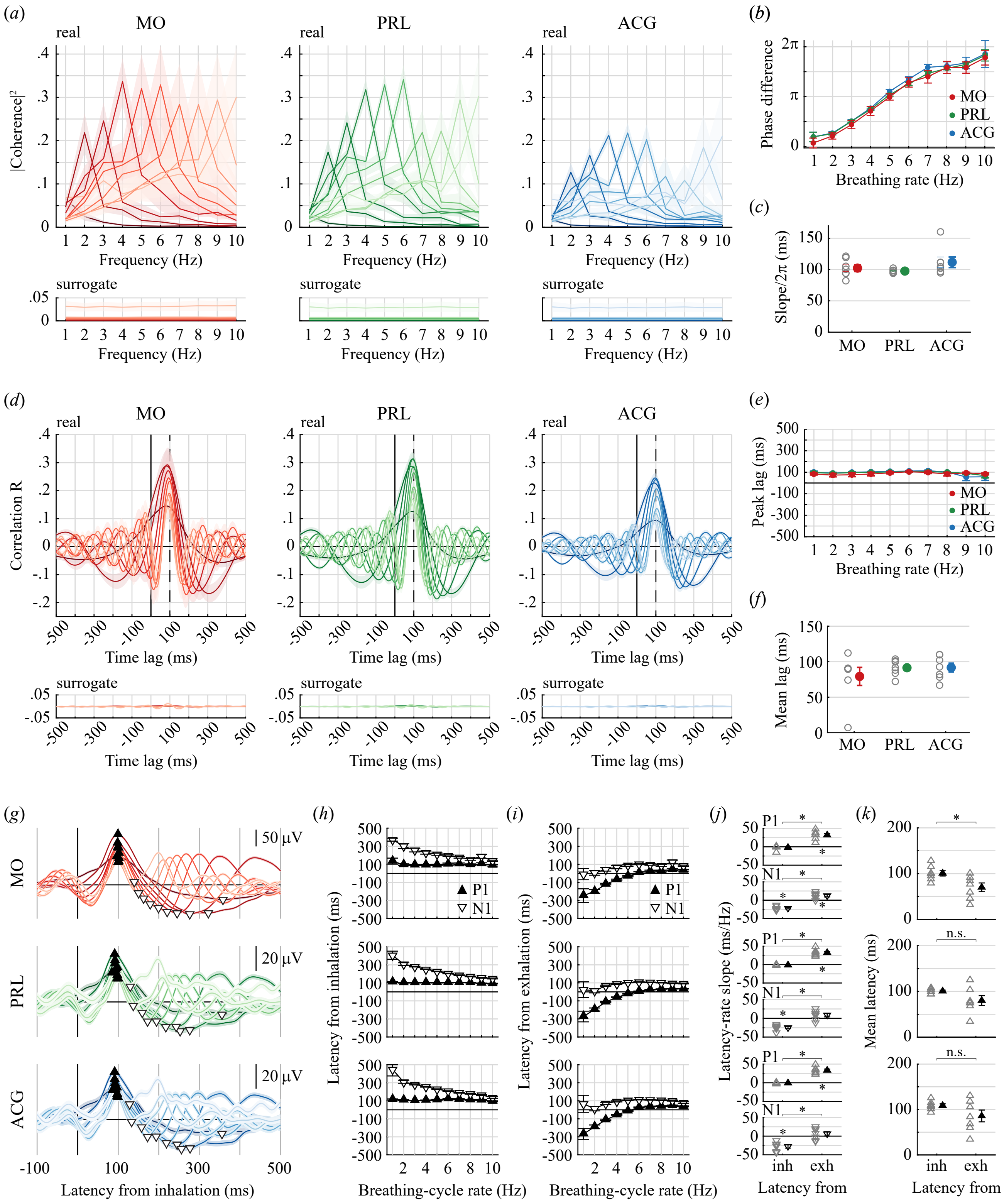


**Figure S1. Alignment of LFPs to nasal breathing across PFC areas.**

**(*a*)** Magnitude-squared-coherence spectra between nasal pressure and average LFPs from the MO, PRL and ACG electrodes, as in Figure 1g. Breathing-brain coherence was significant ($p < 0.01$) for 190 out of 210 combinations of rat, cortical area and breathing rate (Table S1). **(*b*)** Phase of the breathing-brain coherence as a function of breathing rate, as in Figure 1h. A linear regression model revealed a significant effect of breathing rate on phase (LMM $R^2 = 0.85$; slope = $0.195 \pm 0.006\ \pi \cdot \mathrm{Hz}^{-1}$; $F(1,184) = 290.25$, $p < 1e^{-6}$), with significant effect of cortical area on phase at midpoint rate ($F(2,184) = 4.88$, $p = 0.009$; PRL − MO = $-0.13 \pm 0.04\ \pi$, $p = 0.006$) and no interaction between rate and cortical area ($F(2,184) = 0.91$, $p = 0.40$). **(*c*)** Predicted time-lags from nasal pressure to LFPs, as in Figure 1i. No significant effect of cortical area (MO: 102.3 ± 5.0 ms, PRL: 97.6 ± 1.0 ms, ACG: 111.6 ± 7.8 ms; LMM $F(2,18) = 1.51$, $p = 0.25$). ***(d)*** Cross-correlograms between nasal pressure and average LFPs from the MO, PRL and ACG electrodes, as in Figure 2a. Peak correlation R was significant ($p < 0.01$) for 202 of 210 combinations of rat, cortical area and breathing rate (Table S1). ***(e)*** Lag at the LFP-breathing cross-correlation peaks as a function of breathing rate, as in Figure 2b. A linear regression model revealed no effect of breathing rate on peak lag (LMM $R^2 = 0.13$; slope = −0.89 ± 0.67 ms · $Hz^{-1}$; $F(1,196) = 1.44$, $p = 0.23$). The overall model estimate for lag at midpoint rate was 88.7 ± 4.3 ms, with no effect of cortical area ($F(2,196) = 1.94$, $p = 0.15$). There was significant overall interaction between rate and cortical area ($F(2,196) = 3.46$, $p = 0.033$; slope ACG − MO = −4.34 ± 1.66 ms · $Hz^{-1}$; $p = 0.029$). ***(f)*** Means over breathing rates of cross-correlation peak lags, as in Figure 2c (MO: 79.3 ± 12.7 ms, PRL: 91.5 ± 4.2 ms, ACG: 91.7 ± 6.3 ms); no significant difference between cortical areas (LMM; $F(2,18) = 0.69$, $p = 0.51$). ***(g)*** ITAs of MO, PRL and ACG average LFPs for breathing cycles in 1-Hz rate bins, as in Figure 2f. RMS at 0–500 ms latency was significant at $p < 0.01$ for 196 out of 210 combinations of rat, brain region and breathing rate (Table S1). ***(h)*** Latency of P1 and N1 measured from inhalation peaks as a function of breathing-cycle rate, as in Figure 2g. No effect of breathing-cycle rate on P1 latency (LMM $R^2 = 0.097$; slope = −1.67 ± 0.93 ms · $Hz^{-1}$; $F(1,204) = 3.24$, $p = 0.074$), with overall latency at midpoint rate 103.4 ± 3.2 ms and no effect of cortical area ($F(2,204) = 2.93$, $p = 0.055$) or rate-area interaction ($F(2,204) = 0.35$, $p = 0.70$). Significant effect of breathing-cycle rate on N1 latencies (LMM $R^2 = 0.63$; slope = −21.8 ± 2.3 ms · $Hz^{-1}$; $F(1,204) = 90.80$, $p < 1e^{-6}$), with no effect of cortical area on latency at midpoint rate ($F(2,204) = 1.49$, $p = 0.23$) or interaction between rate and area ($F(2,204) = 1.71$, $p = 0.18$). ***(i)*** Latency of P1 and N1 measured from exhalation peaks as a function of breathing-cycle rate, as in Figure 2h. Significant effect of breathing-cycle rate on P1 latencies (LMM $R^2 = 0.65$; slope = 32.1 ± 2.9 ms · $Hz^{-1}$; $F(1,204) = 125.29$, $p < 1e^{-6}$), with no effect of cortical area on latency at midpoint rate ($F(2,204) = 0.31$, $p = 0.74$) or interaction between rate and area ($F(2,204) = 0.037$, $p = 0.96$). Smaller significant effect of breathing-cycle rate on N1 latency (LMM $R^2 = 0.10$; slope = 12.0 ± 3.1 ms · $Hz^{-1}$; $F(1,204) = 15.29$, $p = 0.00013$), with overall latency at midpoint rate 78.7 ± 5.1 ms and no effect of cortical area ($F(2,204) = 0.82$, $p = 0.44$) or rate-area interaction ($F(2,204) = 0.94$, $p = 0.39$). ***(j)*** Linear slopes of P1 and N1 latencies when aligned to times of peak inhalation vs. exhalation, as in Figure 2i. For all cortical areas, slopes of P1 latency vs. rate not significantly different from 0 when measured from inhalations (one-sample t-tests; MO: $t(6) = −0.74$, $p = 0.49$; PRL: $t(6) = −2.12$, $p = 0.078$; ACG: $t(6) = −0.84$, $p = 0.43$). Slopes significantly different from 0 when measuring P1 latency to exhalations (one-sample t-tests; MO: $t(6) = 7.18$, $p = 0.00037$; PRL: $t(6) = 9.59$, $p = 0.000074$; ACG: $t(6) = 9.78$, $p = 0.000066$). Slopes were larger when aligning P1 to exhalations than to inhalations (paired t-tests; $t(6) = 11.32$, $p = 0.000028$, same for all areas). Slopes of N1 latency vs. rate not significantly different from 0 when measured from exhalations for PRL and ACG and significantly different for MO (one-sample t-tests; MO: $t(6) = 3.84$, $p = 0.009$; PRL: $t(6) = 1.82$, $p = 0.12$; ACG: $t(6) = 1.04$, $p = 0.34$). Slopes significantly different from 0 when measuring N1 latency to inhalations (one-sample t-tests; MO: $t(6) = −10.77$, $p = 0.000038$; PRL: $t(6) = −9.21$, $p = 0.000092$; ACG: $t(6) = −6.50$, $p = 0.00063$). Slopes were larger when aligning N1 to inhalations than to exhalations (paired t-tests; $t(6) = 11.32$, $p = 0.000028$, same for all areas). ***(k)*** Means over breathing-cycle rates of latencies of P1 from inhalations and N1 from exhalations, as in Figure 2j. Significant difference for MO only (paired t-tests; MO: $t(6) = −3.70$, $p = 0.010$; PRL: $t(6) = −2.15$, $p = 0.075$; ACG: $t(6) = −2.30$, $p = 0.060$).

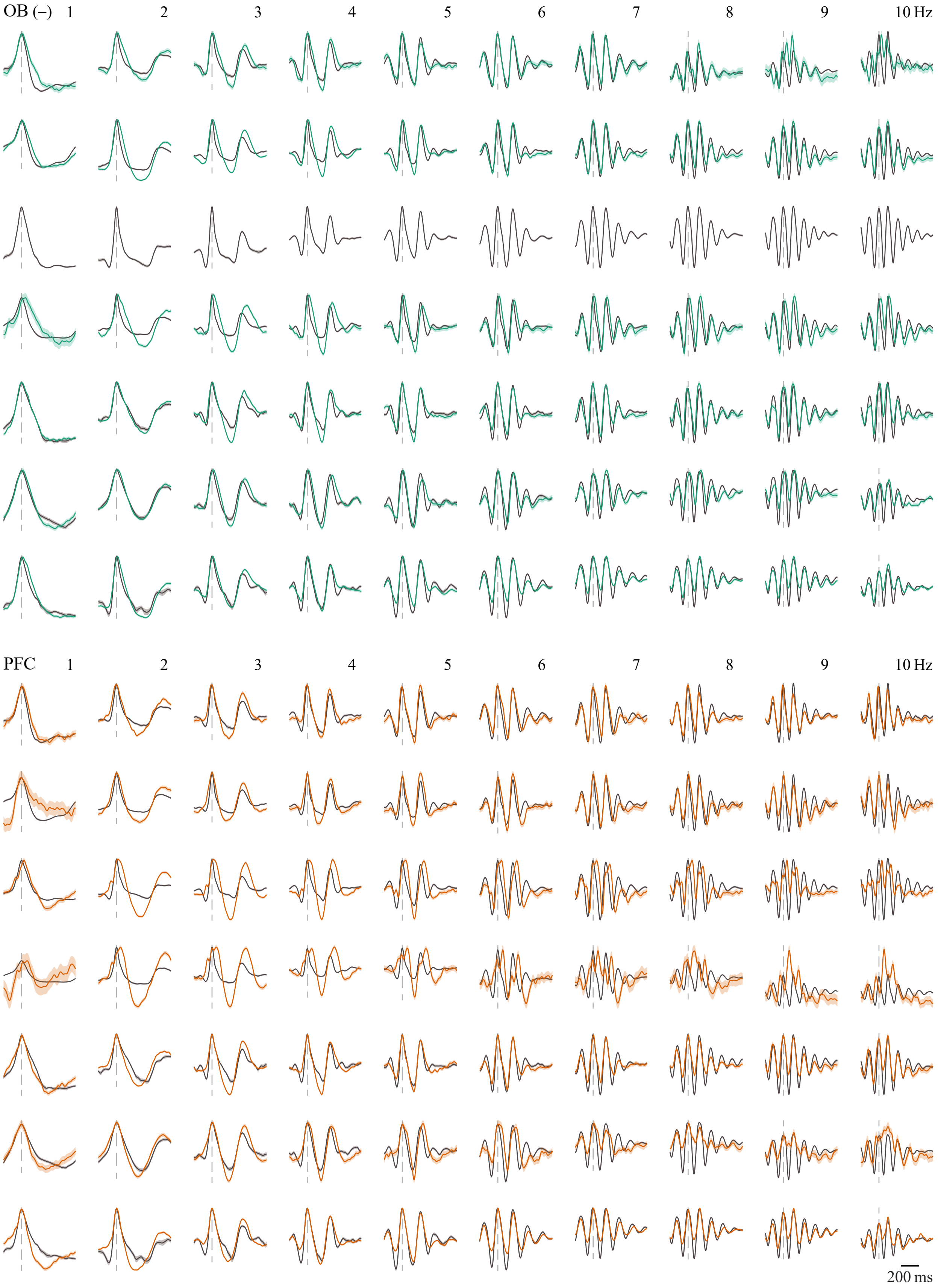


**Figure S2. Average LFP waveforms aligned to inhalations across rats and breathing rates.** Superimposed scaled ITAs of nasal pressure (delayed by 100 ms) and OB or PFC LFPs, for breathing-cycle rate bins of 1 to 10 Hz, as in Figure 2k. One row per rat (OB missing for rat in row #3).

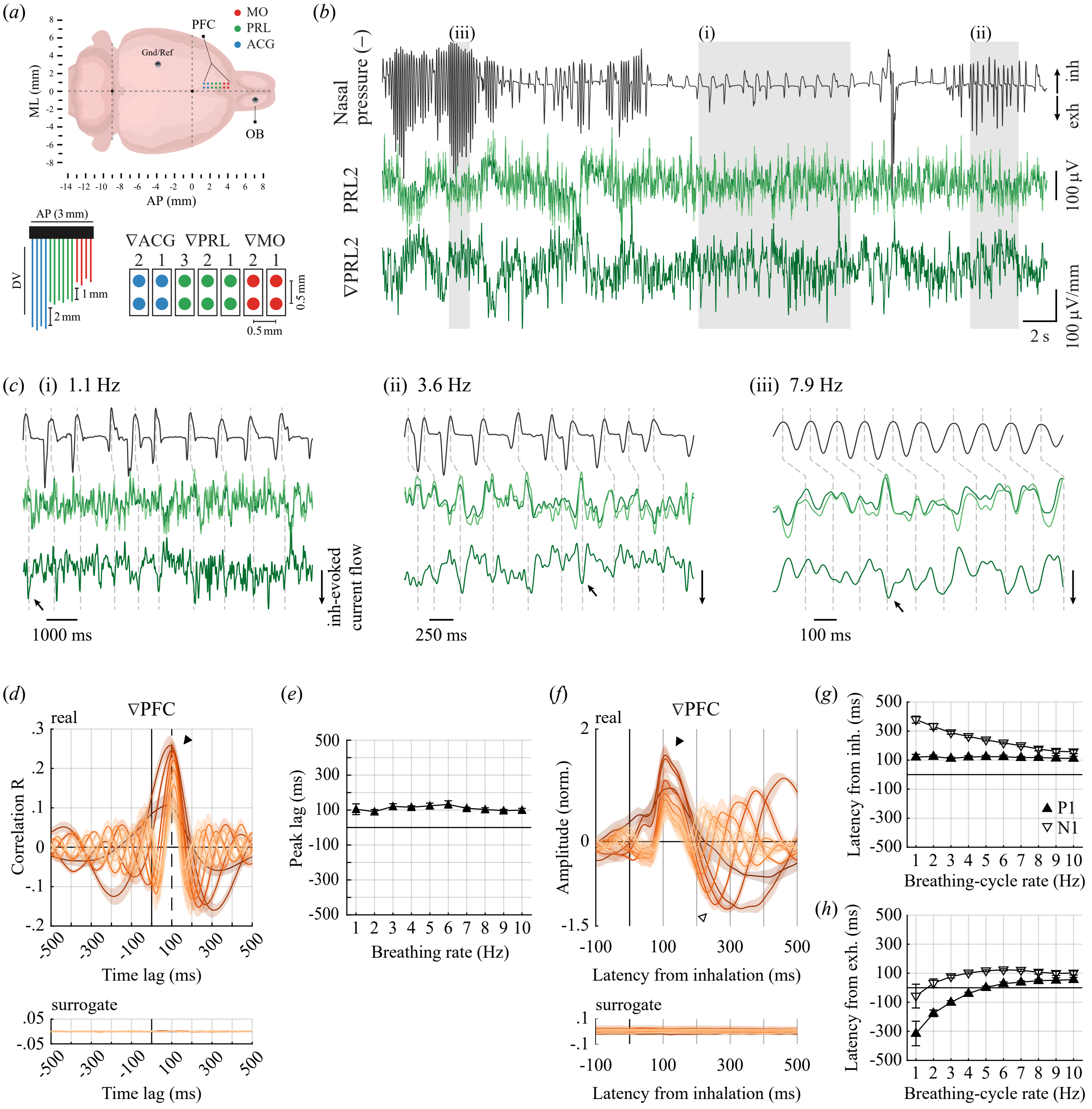


**Figure S3. Time-locking of cortical current flow (∇LFP) to the nasal breathing cycle.**
**(*a*)** Position and geometry of the cortical electrode array. Seven pairs of electrodes extended antero-posteriorly from 4.2 to 1.2 mm from bregma, in two parallel rows positioned at 0.4 and 0.9 mm medio-laterally. We obtained differential LFPs (∇LFP; current flow) by subtracting for each pair (2 in the MO, 3 in the PRL and 2 in the ACG) the signal in the lateral electrode from that of the medial one; positive ∇LFP reflects positive current flow from the medial to the lateral electrode. **(*b*)** Example traces from the same rat and time period in Figure 1b, showing intranasal pressure (.1–20 Hz; top), LFPs (.1–20 Hz) from the medial and lateral electrodes in the PRL2 pair (dark and light green, middle), and ∇LFP between them (bottom). Grayed rectangles mark the time periods expanded in (*c*). **(*c*)** Detailed views of the time periods highlighted in (*b*) for the same signals. Dashed lines connect the times of peak inhalation with 100-ms later in the LFPs for reference. Note negative peaks in current flow density aligning with the 100-ms markers (arrows). We detected those electrode pairs for which the differentiation inverted the polarity of the respiratory modulation and inverted back the ∇LFP signal for all analyses (see Supplementary Methods). **(*d*)** Cross-correlograms between ∇PFC and nasal pressure

across 1–10 Hz breathing rates (mean ± sem over rats; n = 6). ∇PFC is the ∇LFP from the cortical electrode pair with the largest absolute peak correlation R for 2-Hz breathing (4 from MO, 1 from PRL and 1 from ACG). Peak correlation R was significant ($p < 0.01$) for all 60 combinations of rat and breathing rate for ∇PFC, and for 164 out of 170 combinations of rat, cortical area and breathing rate (Table S2). Note how correlations peak at ~100 ms across all breathing rates (filled arrowhead; compare to monopolar LFPs in Figure 2a). (***e***) Lag of the PFC ∇LFPs-breathing cross-correlation peaks as a function of breathing rate. A linear regression model revealed no effect of breathing rate on peak lag (LMM $R^2 = -0.015$; slope = −0.63 ± 1.67 ms · $Hz^{-1}$; $F(1,58) = 0.14$, $p = 0.71$; lag at midpoint rate = 109.9 ± 4.8 ms; compare to monopolar LFPs in Figure 2b). (***f***) ITAs of ∇PFC for breathing cycles in 1–10 Hz rate bins (mean ± sem over rats; n = 6). ∇PFC is the ∇LFP from the cortical electrode pair with the largest RMS (0–500 ms window) in its ITA for 2-Hz breathing cycles (4 from MO, 2 from PRL). The waveforms from each rat were normalized by its 2-Hz RMS value before averaging. Modulation RMS was significant at $p < 0.01$ for 57 out of 60 combinations of rat and breathing-cycle rate for ∇PFC, and for 125 out of 170 combinations of rat, cortical area and breathing-cycle rate (Table S2). Note how waveforms reach a positive peak at ~100 ms across all breathing rates (P1, filled arrowhead), followed by a negative peak whose latency decreases with rate (N1, open arrowhead; compare to example monopolar LFPs in Figure 2f). (***g***) Latency of P1 and N1 measured from inhalation peaks as a function of breathing-cycle rate. No effect of breathing-cycle rate on P1 latency (LMM $R^2 = 0.61$; slope = −1.02 ± 0.76 ms · $Hz^{-1}$; $F(1,54) = 1.81$, $p = 0.18$), with overall latency at midpoint rate 119.4 ± 3.2 ms. Significant effect of breathing-cycle rate on N1 latencies (LMM $R^2 = 0.91$; slope = −23.6 ± 1.13 ms · $Hz^{-1}$; $F(1,54) = 439.21$, $p < 1e^{-6}$). Thus, P1 peaks in cortical current flow follow inhalations with constant latency across breathing-cycle rates (compare to monopolar LFPs in Figure 2g). (***h***) Latency of P1 and N1 measured from exhalation peaks as a function of breathing-cycle rate. Significant effect of breathing-cycle rate on P1 latencies (LMM $R^2 = 0.64$; slope = 34.6 ± 3.4 ms · $Hz^{-1}$; $F(1,54) = 100.77$, $p < 1e^{-6}$). Smaller significant effect of breathing-cycle rate on N1 latencies (LMM $R^2 = 0.31$; slope = 11.7 ± 2.8 ms · $Hz^{-1}$; $F(1,54) = 18.05$, $p = 0.00009$), with overall latency at midpoint rate 86.8 ± 12.4 ms (negative N1 latencies for 1-Hz breathing appear partly driven by noisy peak detection; overall mean latency excluding the 1-Hz bin was 97.8 ± 8.6 ms). Thus, N1 peaks in cortical current flow follow exhalations with a nearly constant latency across breathing-cycle rates (compare to monopolar LFPs in Figure 2h). One rat had only one valid electrode pair, located in the ACG, with no significant respiratory modulation throughout and was excluded from all analyses of ∇LFP.

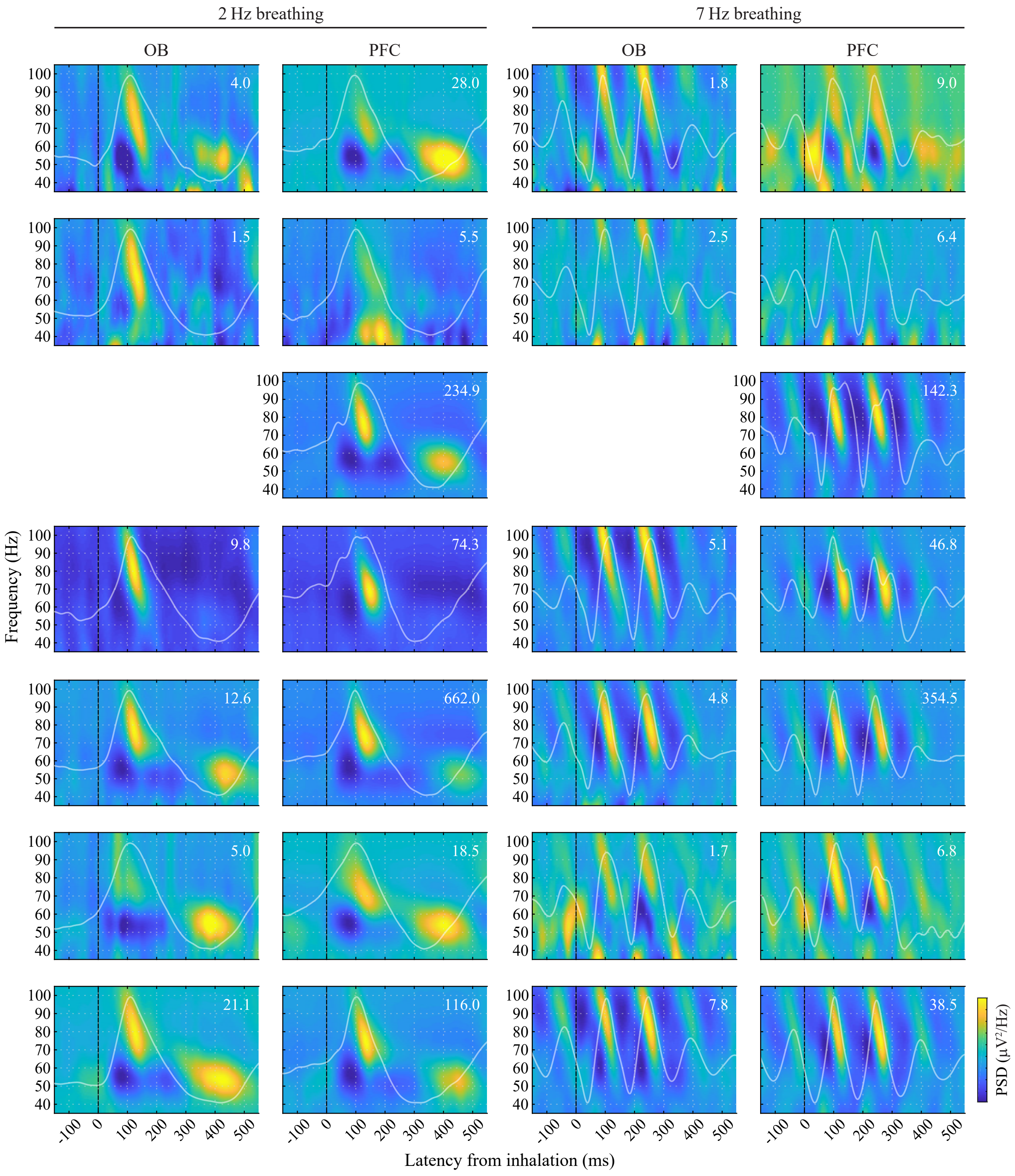


**Figure S4. Average gamma spectrogram aligned to inhalations across rats.**
Gamma-spectrogram ITA for breathing cycles in the 2- and 7-Hz rate bins, for the OB and PFC electrodes of each rat, as in Figure 3c, d (one row per rat). Each panel individually scaled, with the range of power indicated at the top-right for each. Slow gamma was not evident for rats in rows #2 and 4 and was thus excluded from all analyses.

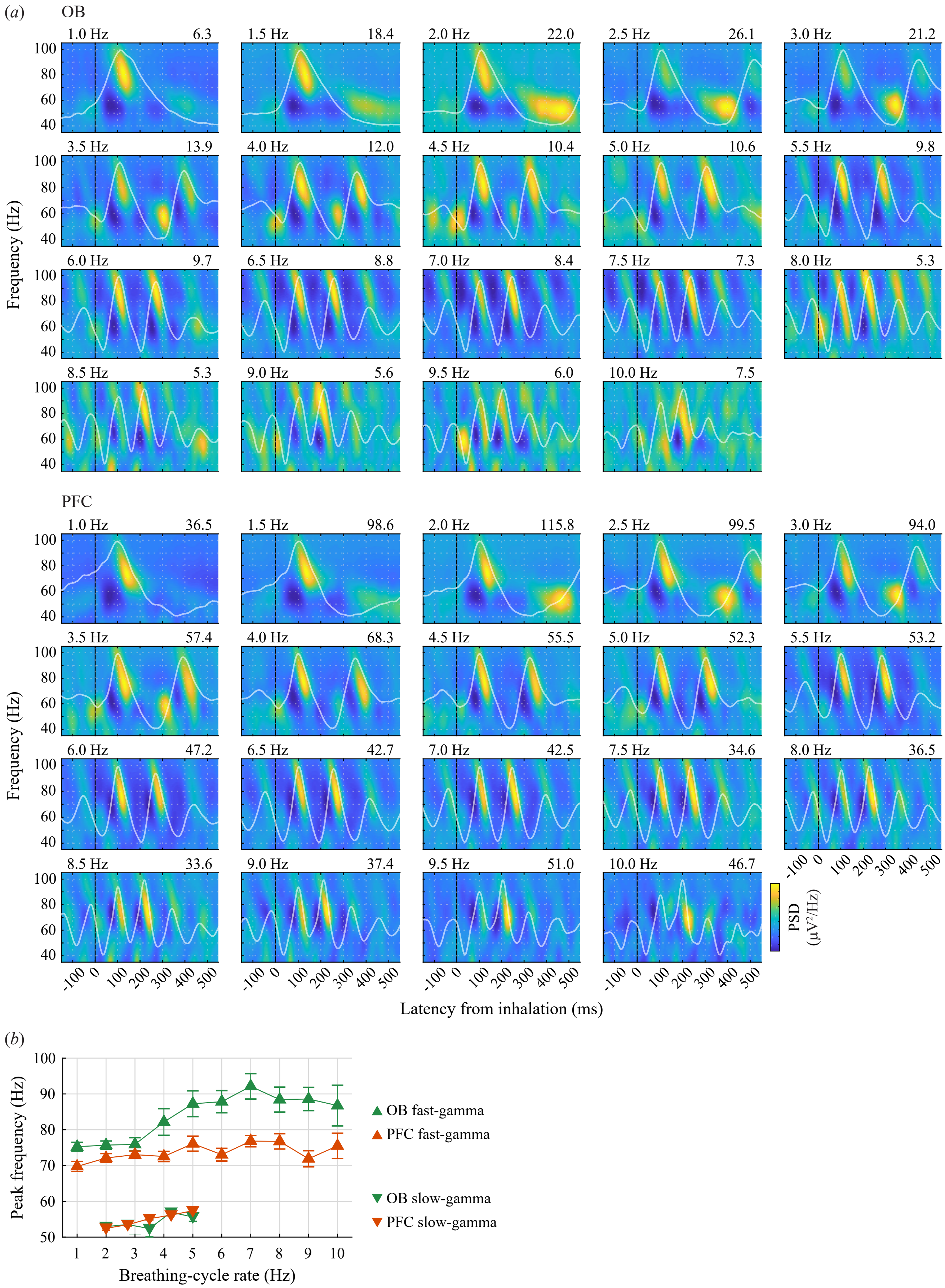


**Figure S5. Respiratory fast- and slow-gamma bursts across breathing-cycle rates.**

**(*a*)** Detailed view of the gamma-spectrogram ITA across breathing-cycle rates for the OB and PFC electrodes from one example rat, as in Figure 3c, d. Each panel individually scaled, with the range of power indicated at the top-right for each. Note how fast-gamma bursts are visible for all rates, while slow-gamma mostly for ~1.5 to 4 Hz. ***(b)*** Peak frequency of fast- and slow-gamma bursts as a function of breathing-cycle rates (mean ± sem over rats). Peaks were calculated within lag-frequency boundaries chosen to capture bursts across all rats and rates (fast gamma: 80–170 ms from inhalation peak, 65–105 Hz; slow gamma: 50–200 ms from exhalation peak, 45–60 Hz). Peak frequency of fast gamma increased with breathing-cycle rate (LMM $R^2 = 0.51$; slope = 1.11 ± 0.20; $F(1,126) = 36.11$, $p < 1e^{-6}$; n = 6 and 7 rats for OB and PFC) and its frequency at midpoint rate and slope were lower for PFC (frequency: PFC − OB = −10.6 ± 1.2 Hz; $F(1,126) = 84.44$, $p < 1e^{-6}$; slope: PFC − OB = −1.26 ± 0.39; $F(1,126) = 10.24$, $p = 0.0017$).

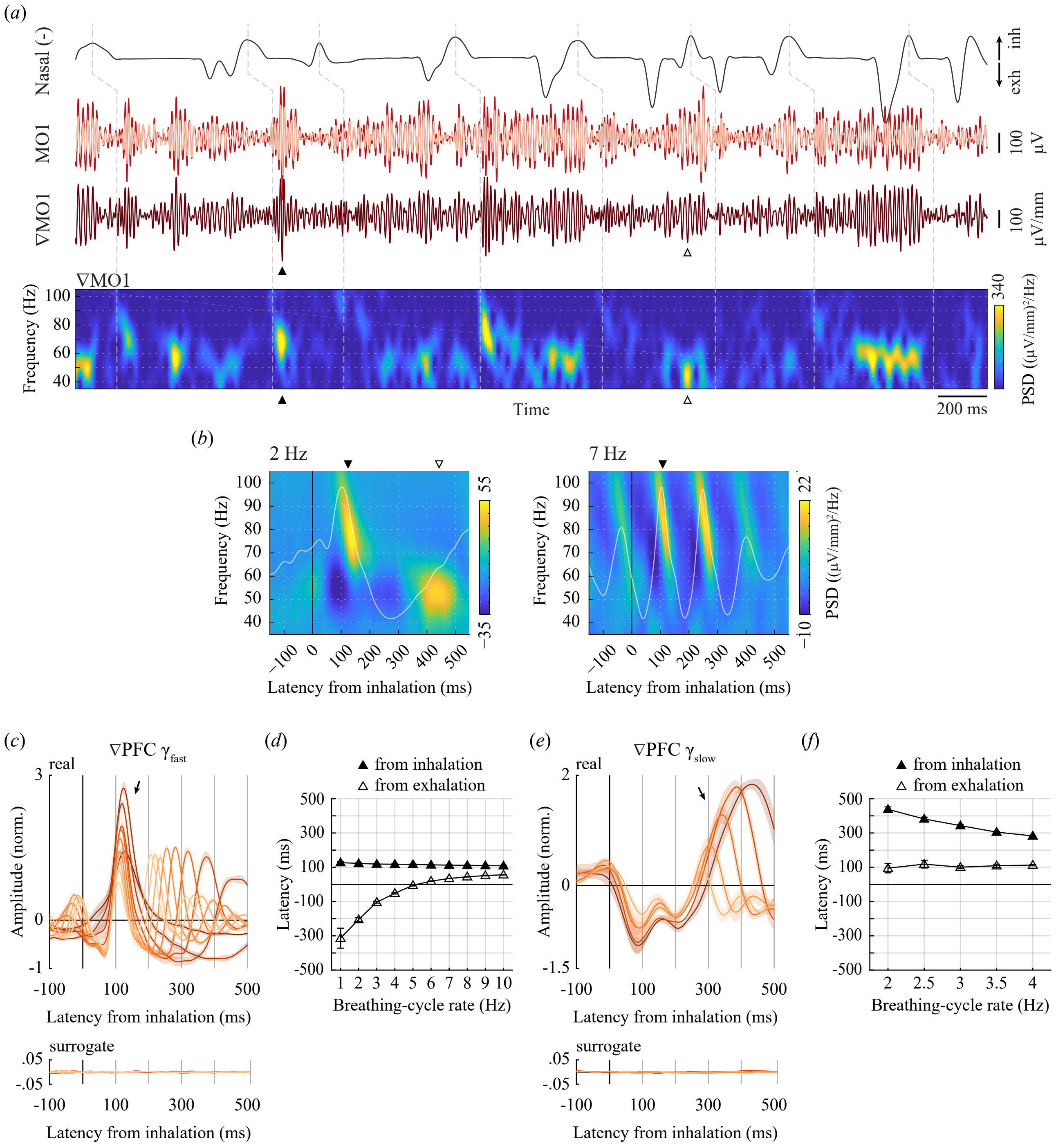


**Figure S6. Time-locking of fast- and slow-gamma bursts of cortical current flow (∇LFP) to the nasal breathing cycle.**

**(a)** Top: Example traces from the same rat and time period in Figure 3a (zoomed in, mean breathing-cycle rate 2.7 Hz), showing intranasal pressure (.1–20 Hz; top), gamma LFPs (40–100 Hz) from the medial and lateral electrodes in the MO1 pair (dark and light traces, middle) and current flow (∇LFP) between them (darker trace, bottom). Bottom: spectrogram of the ∇LFP. Dashed lines connect the times of peak inhalation with 100 ms later in the LFPs for reference. Note how salient fast- and slow-gamma bursts in cortical current flow (closed and open arrowheads) align with inhalations and exhalations. ***(b)*** ITA of the ∇MO1 gamma spectrogram for all breathing cycles in the 2 and 7 Hz rate bins for the same rat in (*a*). The waveforms of the ∇MO1 LFP ITAs are superimposed in white. Compare to their monopolar counterparts in Figure 3c, d. ***(c)*** ITAs of the ∇PFC fast-gamma envelope for breathing cycles in 1–10 Hz rate bins (1-Hz bins; mean ± sem over 5 rats with significant modulations). ∇PFC is the ∇LFP from the cortical electrode pair with the largest RMS (0–200 ms window) in its fast-gamma envelope ITA for 2-Hz breathing cycles (4 from MO, 1 from PRL). The waveforms from each rat were normalized by their 2-Hz RMS value before averaging (mean RMS = 12.4 μV/mm, range 2.7–21.0). Modulation RMS was significant at $p < 0.01$ for 50 out of 60 combinations of rat and breathing-cycle rate for ∇PFC, and for 85 out of 170 combinations of rat, cortical area and breathing-cycle rate (Table S2). Note how fast-gamma envelopes reach a positive peak shortly after 100 ms across all rates (arrow; compare to example monopolar fast-gamma envelopes in Figure 3e). **(*d*)** Latency of ∇PFC fast-gamma amplitude peaks from inhalation or exhalation peaks as a function of breathing-cycle rate. Latency from inhalation was subtly but significantly dependent on rate (LMM $R^2 = 0.83$; slope = −1.88 ± 0.16 ms · $Hz^{-1}$; $F(1,48) = 133.01$, $p < 1e^{-6}$), with overall latency at midpoint rate 115.8 ± 2.35 ms. Latency from exhalation was pronouncedly dependent on rate (LMM $R^2 = 0.73$; slope = 37.3 ± 3.2 ms · $Hz^{-1}$; $F(1,48) = 135.6$, $p < 1e^{-6}$). Thus, fast-gamma bursts in cortical current flow follow inhalations with close-to-constant latency across breathing-cycle rates (compare to monopolar in Figure 3f). **(*e*)** ITAs of the ∇PFC slow-gamma envelope for breathing cycles in 2–4 Hz rate bins (0.5-Hz bins; mean ± sem over 4 rats with significant modulations). ∇PFC is the ∇LFP from the cortical electrode pair with the largest RMS (0–200 ms window, from exhalation) in its slow-gamma envelope ITA for 2-Hz breathing cycles (3 from MO, 1 from PRL). The waveforms from each rat were normalized by their 2-Hz RMS value before averaging (mean RMS = 8.43 μV/mm, range 2.3–17.6). Modulation RMS was significant at $p < 0.01$ for 20 out of 20 combinations of rat and breathing-cycle rate for ∇PFC, and for 28 out of 60 combinations of rat, cortical area and breathing-cycle rate (Table S2). Note how slow-gamma envelopes reach a positive peak at a latency that decreases with rate (arrow; compare to example monopolar slow-gamma envelopes in Figure 3g). **(*f*)** Latency of ∇PFC slow-gamma amplitude peaks from inhalation or exhalation peaks as a function of breathing-cycle rate. Latency from inhalation was pronouncedly dependent on rate (LMM $R^2 = 0.95$; slope = −77.4 ± 4.3 ms · $Hz^{-1}$; $F(1,18) = 326.02$, $p < 1e^{-6}$). Latency from exhalation was not significantly dependent on rate (LMM $R^2 = 0.31$; slope = 5.16 ± 8.0 ms · $Hz^{-1}$; $F(1,18) = 0.42$, $p = 0.53$), with overall latency at midpoint rate (3 Hz) 107.0 ± 11.9 ms. Thus, slow-gamma bursts in cortical current flow follow exhalations with constant latency across breathing-cycle rates (compare to monopolar in Figure 3h).

| Metric | Brain region | Breathing rate | | | | | | | | | |
|---|---|---|---|---|---|---|---|---|---|---|---|
| | | 1 Hz | 2 Hz | 3 Hz | 4 Hz | 5 Hz | 6 Hz | 7 Hz | 8 Hz | 9 Hz | 10 Hz |
| **Breathing-brain coherence magnitude (Figure 1g–i and S1a–c)** | **OB (6)** | 6 | 6 | 6 | 6 | 6 | 6 | 6 | 6 | 5 | 5 |
| | **PFC (7)** | 5 | 7 | 7 | 7 | 7 | 7 | 6 | 6 | 7 | 6 |
| | **MO (7)** | 7 | 7 | 7 | 7 | 7 | 6 | 6 | 5 | 6 | 4 |
| | **PRL (7)** | 6 | 7 | 7 | 7 | 7 | 7 | 7 | 7 | 7 | 5 |
| | **ACG (7)** | 6 | 7 | 7 | 7 | 7 | 7 | 7 | 6 | 5 | 2 |
| **Cross-correlation peak R (Figure 2a–c and S1d–f)** | **OB (6)** | 6 | 6 | 6 | 6 | 6 | 6 | 6 | 6 | 6 | 6 |
| | **PFC (7)** | 7 | 7 | 7 | 7 | 7 | 7 | 7 | 7 | 7 | 7 |
| | **MO (7)** | 6 | 7 | 7 | 7 | 7 | 6 | 7 | 7 | 6 | 6 |
| | **PRL (7)** | 7 | 7 | 7 | 7 | 7 | 7 | 7 | 7 | 7 | 6 |
| | **ACG (7)** | 7 | 7 | 7 | 7 | 7 | 7 | 7 | 7 | 7 | 4 |
| **LFP ITA RMS (0–500 ms) (Figure 2f–k and S1g–k)** | **OB (6)** | 6 | 6 | 6 | 6 | 6 | 6 | 6 | 6 | 6 | 6 |
| | **PFC (7)** | 6 | 7 | 7 | 7 | 7 | 7 | 7 | 7 | 7 | 7 |
| | **MO (7)** | 5 | 6 | 6 | 6 | 6 | 6 | 6 | 6 | 6 | 6 |
| | **PRL (7)** | 6 | 7 | 7 | 7 | 7 | 7 | 7 | 7 | 7 | 7 |
| | **ACG (7)** | 5 | 7 | 7 | 7 | 7 | 7 | 7 | 7 | 7 | 7 |
| **Fast-gamma envelope ITA RMS (0–200 ms) (Figure 3e, f)** | **OB (6)** | 6 | 6 | 6 | 6 | 6 | 5 | 4 | 3 | 4 | 5 |
| | **PFC (7)** | 6 | 7 | 7 | 7 | 7 | 6 | 6 | 6 | 6 | 6 |
| | **MO (7)** | 6 | 6 | 6 | 6 | 6 | 6 | 6 | 6 | 6 | 6 |
| | **PRL (7)** | 5 | 7 | 7 | 7 | 6 | 6 | 6 | 6 | 4 | 5 |
| | **ACG (7)** | 6 | 6 | 6 | 6 | 6 | 6 | 5 | 4 | 3 | 3 |
| **Slow-gamma envelope ITA RMS (0–200 ms from exhalation) (Figure 3g, h)** | **OB (4)** | - | 4 | 4 | 3 | - | - | - | - | - | - |
| | **PFC (5)** | - | 5 | 5 | 5 | - | - | - | - | - | - |
| | **MO (5)** | - | 5 | 5 | 4 | - | - | - | - | - | - |
| | **PRL (5)** | - | 5 | 5 | 2 | - | - | - | - | - | - |
| | **ACG (5)** | - | 5 | 5 | 1 | - | - | - | - | - | - |

**Table S1. Statistical significance of metrics per subject, brain region and breathing rate.** Each cell represents the number of rats for which a given metric was significant for each breathing-rate bin ($\alpha = 0.01$). For coherence and cross-correlation, breathing rate is the frequency with peak power in the nasal-pressure PSD for 1-second windows. For ITAs, it is the instantaneous rate of each breathing cycle. PFC is the cortical channel with largest breathing modulation for each rat. For coherence, cross-correlation and LFP ITA, a rat counts as significant for MO, PRL or ACG if a metric was significant for the average LFP from that area. For the more local fast and slow gamma, a rat counts as significant for MO, PRL or ACG if any of its valid electrodes within that area were significant, with p-values corrected with Holm-Bonferroni within each pool of 1 to 6 valid electrodes. Numbers in parentheses are the number of rats with valid data for each area. For slow gamma, the 2 rats with no visible modulation are excluded from the count. Values for slow gamma are from 0.5-Hz bins (2.5 and 3.5 Hz not shown), all others from 1-Hz bins.

| Metric | Brain region | Breathing rate | | | | | | | | | |
|---|---|---|---|---|---|---|---|---|---|---|---|
| | | 1 Hz | 2 Hz | 3 Hz | 4 Hz | 5 Hz | 6 Hz | 7 Hz | 8 Hz | 9 Hz | 10 Hz |
| ∇LFP Cross-correlation peak R (Figure S3d, e) | PFC (6) | 6 | 6 | 6 | 6 | 6 | 6 | 6 | 6 | 6 | 6 |
| | MO (6) | 5 | 6 | 6 | 6 | 6 | 6 | 6 | 6 | 6 | 6 |
| | PRL (6) | 4 | 6 | 6 | 6 | 6 | 6 | 6 | 6 | 6 | 6 |
| | ACG (5) | 3 | 4 | 5 | 5 | 5 | 5 | 5 | 5 | 5 | 5 |
| ∇LFP ITA RMS (0–500 ms) (Figure S3f, g) | PFC (6) | 4 | 6 | 6 | 6 | 6 | 6 | 6 | 6 | 5 | 6 |
| | MO (6) | 3 | 4 | 4 | 5 | 5 | 5 | 5 | 5 | 5 | 5 |
| | PRL (6) | 4 | 4 | 6 | 5 | 5 | 5 | 5 | 4 | 4 | 5 |
| | ACG (5) | 3 | 4 | 4 | 4 | 3 | 3 | 3 | 2 | 3 | 3 |
| ∇LFP fast-gamma envelope ITA RMS (0–200 ms) (Figure S6c, d) | PFC (6) | 5 | 5 | 5 | 5 | 5 | 5 | 5 | 5 | 5 | 5 |
| | MO (6) | 5 | 5 | 5 | 5 | 5 | 5 | 5 | 5 | 5 | 5 |
| | PRL (6) | 2 | 4 | 4 | 3 | 2 | 2 | 1 | 1 | 1 | 2 |
| | ACG (5) | 2 | 3 | 3 | 1 | 1 | 1 | 1 | 0 | 0 | 1 |
| ∇LFP slow-gamma envelope ITA RMS (0–200 ms from exhalation) (Figure S6e, f) | PFC (4) | - | 4 | 4 | 4 | - | - | - | - | - | - |
| | MO (4) | - | 4 | 4 | 3 | - | - | - | - | - | - |
| | PRL (4) | - | 1 | 1 | 1 | - | - | - | - | - | - |
| | ACG (4) | - | 2 | 1 | 0 | - | - | - | - | - | - |

**Table S2. Statistical significance of metrics for cortical current flow (∇LFP) per subject, brain region and breathing rate.** Each cell represents the number of rats for which a given metric was significant for each breathing-rate bin ($\alpha = 0.01$). PFC is the ∇LFP with the largest breathing modulation over all valid electrode pairs for each rat. For MO, PRL and ACG, a rat counts as significant if any of its valid electrode pairs within that area were significant, with p-values corrected with Holm-Bonferroni within each pool of 1 to 3 valid pairs. Numbers in parentheses are the number of rats with valid ∇LFP data for each area. For slow gamma, the 2 rats with no visible modulation are excluded from the count. One rat had only one valid pair, located in the ACG, with no significant respiratory modulation throughout and was excluded from all analyses. Values for slow gamma are from 0.5-Hz bins (2.5 and 3.5 Hz not shown), all others from 1-Hz bins.

### Supplementary Methods

#### Current flow density

For the analysis of current flow density (∇LFP; Figures S3 and S6), we obtained differential LFPs between pairs of cortical electrodes (0.5 mm distance; 2 pairs in the MO, 3 in the PRL and 2 in the ACG; Figure S3a) by subtracting the signals in the lateral electrode from that of the medial one and divided the differential signals by the electrode distance (Figure S3b); positive ∇LFP reflects positive current flow from the medial to the lateral electrode. We note that ∇LFP is not in units of current as it was not scaled by an estimate of the tissue conductivity [1]. Only pairs where both electrodes had valid signals were included from each rat. One rat had only one valid electrode pair, located in the ACG, with no significant respiratory modulation throughout and was excluded from all analyses of ∇LFP.

Similar to the analyses of monopolar PFC signals, we selected for ∇LFP the electrode pair with the most salient breathing modulation at 2-Hz breathing as representative of PFC for each rat on a per-analysis basis (named ∇PFC), as detailed in the legends of Figures S3 and S6. For some electrode pairs, the differentiation inverted the sign of the breathing-modulated signals (e.g. Figure S3b). For the cross-correlation and ITA analyses in Figure S3d–h we detected and inverted back these cases before computing latencies and averaging waveforms across rats. In detail, for the cross-correlation between ∇LFP and nasal pressure (Figure S3d), we computed for each electrode pair and breathing-rate bin the Pearson correlation between the cross-correlation function for ∇LFP and the mean of the functions for the 2 monopolar LFPs in the ±0.5 s lag window and identified inversions as cases with negative R across breathing-rate bins (3 out of 6 ∇PFC inverted). For ∇LFP ITAs, we performed an equivalent analysis on the 0–0.5 s window of the ITA waveforms (4 out of 6 ∇PFC inverted; the ∇PFC pair selected was different for cross-correlation and ITA for one rat).


1. Kwan HC, Murphy JT. 1974 A basis for extracellular current density analysis in cerebellar cortex. *J. Neurophysiol.* **37**, 170–180. (doi:10.1152/jn.1974.37.1.170)
